# A ROS-dependent mechanism to drive progression through S phase

**DOI:** 10.1101/2022.03.31.486607

**Authors:** Dilyana Georgieva Kirova, Kristyna Judasova, Julia Vorhauser, Thomas Zerjatke, Jacky Kieran Leung, Ingmar Glauche, Jörg Mansfeld

**Author notes:** These authors contributed equally to this work (Dilyana Georgieva Kirova, Kristyna Judasova).

## Abstract

Long considered as cytotoxic reagents, reactive oxygen species (ROS) at the right concentration promote cell proliferation in cell culture, stem cells and model organisms. However, how ROS signaling is coordinated with cell cycle progression and integrated into the cell cycle control machinery on the molecular level remains unsolved. Here, we report oscillations of mitochondrial ROS during the cell cycle that target cyclin-dependent kinase 2 (CDK2). Chemical and metabolic interference with ROS production decrease T-loop phosphorylation on CDK2, impeding its full activation and thus efficient DNA replication. ROS regulate CDK2 activity through oxidation of a conserved cysteine residue in close proximity to the T-loop, which prevents binding of the T-loop phosphatase KAP. Together our data reveal how ROS couple mitochondrial metabolism to DNA replication and cell cycle progression, and provide a solution to the longstanding conundrum of how KAP activity towards CDKs can be cell cycle-regulated.

## INTRODUCTION

Reactive oxygen species (ROS) are oxygen-containing molecules of high chemical reactivity based on their ability to “steal” electrons from molecules (oxidation). Overwhelming exposure to ROS during oxidative stress or in response to external sources such as ionizing agents oxidize proteins, cause mutations in DNA, and trigger lipid peroxidation (Schieber & Chandel 2014). The major physiological sources of cellular ROS are membrane-associated NADPH oxidases (NOXs) and mitochondria, which produce superoxide anions (O2^•−^) in response to growth factor signaling and as by-product of oxidative phosphorylation, respectively (Schieber & Chandel 2014; Bedard & Krause 2007; Murphy 2009). Short-lived and membrane-impermeant O2^•−^ molecules are subsequently converted to hydrogen peroxide (H_2_O_2_), either spontaneously or catalytically by superoxide dismutases (Möller et al. 2019; Y. Wang et al. 2018). H_2_O_2_ is more stable and membrane-permeant and acts as a physiological signaling molecule in diverse biological processes including growth factor (GF) signaling, proliferation, differentiation, and adaptation to hypoxia (Holmström & Finkel 2014; Reczek & Chandel 2017; Shadel & Horvath 2015). Physiological ROS signaling is in part mediated by reversible oxidation of cysteine residues to sulfenic acids (R-SOH) and intra- or intermolecular disulfide bonds, which act as reversible posttranslational modifications (PTMs) regulating activity, localization, stability and interactions of proteins (Reddie & Carroll 2008).

In recent years low concentrations of ROS, in particular of H_2_O_2_, have been linked to increased proliferation of stem cells, differentiated cells and cancer cells (Armstrong et al. 2010; Gurusamy et al. 2009; Adusumilli et al. 2020; Ahmed Alfar et al. 2017; Moll et al. 2018; Sigaud et al. 2005; Safford et al. 1994; Irani et al. 1997; Ogrunc et al. 2014). Mechanistically, this pro-proliferative effect is best understood for growth factor (GF) signaling, where NOX-derived H_2_O_2_ activates membrane-associated receptor tyrosine kinases while inhibiting counteracting protein tyrosine phosphatases (Holmström & Finkel 2014); consequently, sustained GF signaling initiates transcriptional programs that promote the G0/G1 and G1/S transitions (Burhans & Heintz 2009; Chiu & Dawes 2012). Mitochondrial ROS can also activate transcriptional programs linked to proliferation (Owusu-Ansah et al. 2008; Weinberg et al. 2010; Tsai et al. 2011; Connor et al. 2005); however, whether or not ROS also directly target cell cycle regulation independently of transcriptional signaling remains elusive.

Cell cycle regulation is executed by cyclin-dependent kinases (CDKs), which are activated by a conserved two-step mechanism: binding of a cell cycle stage-specific cyclin and phosphorylation of a threonine residue (T160 in CDK2) within the activation segment of the kinase domain (T-loop) (Morgan 2007). T-loop phosphorylation is carried out by a trimeric CDK-activating kinase (CAK, a trimeric complex of CDK7, cyclin H and MAT1) and opposed by CDK-associated phosphatase (KAP) and a protein phosphatase 2C-like protein (Fesquet et al. 1993; Poon et al. 1993; Solomon et al. 1993; Fisher & Morgan 1994; Mäkelä et al. 1994; Poon & Hunter 1995; Hannon et al. 1994; Gyuris et al. 1993; Cheng et al. 1999). Whether T-loop phosphorylation is regulated to ensure that the right CDK becomes fully active at the right time is still debated, since CAK is considered to be constitutively active (Fisher 2005) and KAP activity regulation is unclear. Cell cycle stage and CDK specificity of T-loop phosphorylation, i.e. on CDK2 and CDK1, is in part achieved by preferential binding of CAK to monomeric CDK2 and cyclin-complexed CDK1 *in vivo*, respectively (Larochelle et al. 2007; Merrick et al. 2008; Fisher & Morgan 1994; Desai et al. 1995). While KAP binds to both, monomeric and cyclin-complexed CDKs (Poon & Hunter 1995; Gyuris et al. 1993; Hannon et al. 1994), it only dephosphorylates monomeric CDKs, i.e. once the bound cyclin is degraded (Poon & Hunter 1995). This results in a conceptual problem for CDK2 activation during S phase, when cyclin E is degraded and CDK2 switches from cyclin E to cyclin A, meaning that T160 can be targeted by CAK and KAP at the same time.

ROS derived from mitochondrial respiration are excellent candidates as direct regulators of the core cell cycle machinery since mitochondrial metabolism, morphology, and activity are tightly coordinated with cell cycle progression (Schieke et al. 2008; Lopez-Mejia & Fajas 2015; Huber et al. 2020). For instance, inactivation of the anaphase promoting complex/cyclosome (APC/C) at the G1/S transition increases the influx of metabolites into the tricarboxylic acid (TCA) cycle (Colombo et al. 2010; Wu et al. 2019) and regulates mitochondrial hyperfusion (Horn et al. 2011), events linked to the entry into S phase (Mitra et al. 2009). In G2 phase, CDK1-driven assembly of the outer mitochondrial membrane translocase and potentially phosphorylation of complex I in the electron transport chain (ETC) increases respiration (Harbauer et al. 2014; Z. Wang et al. 2014). These findings imply the existence of bidirectional feedback between mitochondrial metabolism and cell cycle regulation. However, potential targets of mitochondrial ROS within the core cell cycle machinery and the underlying molecular mechanisms that drive proliferation are yet to be defined.

Here we employ retina pigment epithelial cells (RPE-1) as a non-transformed cell cycle model to monitor and functionally interrogate cellular ROS production in relation to cell cycle progression and CDK2 activation. We uncover periodical oscillations of mitochondrial ROS across the cell cycle that recapitulate the activity profile of S phase-regulating CDK2. We show that oxidation of a conserved cysteine that is only found in CDK2 but no other CDKs ensures T-loop phosphorylation required for efficient DNA replication and progression through S phase. Finally, our data reveal a mechanism for CDK2 activity regulation via KAP that provides positive feedback from mitochondrial metabolism to control the cell cycle and proliferation.

## RESULTS

### Proliferation and S phase progression require oxidative events

To investigate the interplay between physiological ROS signaling and cell cycle progression, we chose hTERT-immortalized RPE-1 cells as a well-established tissue cell culture model with unperturbed cell cycle control (Bodnar et al. 1998). We first assessed whether RPE-1 cells require oxidative events for normal proliferation, as has been reported for other experimental models (Wartenberg et al. 1999; Havens et al. 2006; Murrell et al. 1990; Ohguro et al. 1999; Paul et al. 2014; Ahmed Alfar et al. 2017). Indeed, treating RPE-1 cells with increasing concentrations of the antioxidant N-acetyl-L-cysteine (NAC) for 48 hours reduced proliferation in a dose-dependent manner (Figure S1A). Notably, the decrease in proliferation was not due to induction of apoptosis as monitored by PARP cleavage (Figure S1B). To determine precisely the cell cycle stages that are sensitive to reducing conditions, we employed RPE-1 cells expressing three endogenous proteins tagged with different fluorescent proteins: i) proliferating cell nuclear antigen (Ruby-PCNA) to detect replication foci in S phase, ii) cyclin A2-Venus to distinguish cyclin A2-negative G1 phase cells from cyclin A2-positive S and G2 phase cells, and iii) histone 3.1-turquoise2 (H3.1-Turq2) to segment nuclei (Zerjatke et al. 2017). Combining the features of PCNA and cyclin A2 localization and expression, respectively, allowed us univocally to assign G1, S, and G2 phases from snapshots (Figure S1C). This revealed that at all concentrations the addition of NAC significantly increased the proportion of cells in S phase. In addition, we observed that low (6 mM) concentrations of NAC reduced the number of cells in G1, whereas higher concentrations (8 mM and 10 mM) reduced the number of cells in G2 (Figure S1D). As a general reductant NAC does not distinguish between different forms of ROS, therefore we added membrane-permeant polyethylene glycol-linked catalase (PEG-CAT) to target H_2_O_2_ specifically, which is considered the second messenger of ROS signaling due to its longer half-life and membrane permeability (Winterbourn 2008). To distinguish all cell cycle phases on snapshots, we combined this time Ruby-PCNA and H3.1-Turq2 with Fucci-Gem1-110, a well-established APC/C activity reporter that accumulates after the G1/S transition, when the APC/C is inactivated (Sakaue-Sawano et al. 2008). Fucci-Gem1-110 marks S and G2 phase cells that can subsequently be separated into individual cell cycle phases based on Ruby-PCNA-visualized replication foci (Figure 1A). As with NAC, adding PEG-CAT strongly reduced cell proliferation (Figure 1B) and significantly increased the fraction of cells in S phase at the expense of G1 phase (Figure 1C). Finally, to assess whether ROS directly promotes proliferation in RPE-1 cells, we decided to increase the intracellular levels of ROS. To this end we used genetically encoded D-amino acid oxidase (DAO) from *Rhodosporiduim toruloides*, which converts medium-supplied D-alanine to a keto-acid, ammonia and H_2_O_2_ (Lee & Chu 1996) (Figure 1D). The increase of H_2_O_2_ in response to D- but not L-alanine can be directly monitored by ratio imaging taking advantage of the ratio-metric H_2_O_2_ sensor HyPer2 fused to the N-terminus of DAO (Figure 1E) (Matlashov et al. 2014). DAO was further linked to a nuclear export sequence (NES, HyPer2-DAO-NES) to recapitulate mitochondrial ROS that is released in the form of H_2_O_2_ through the outer mitochondrial membrane (Shadel & Horvath 2015). Indeed, the lowest concentration of D-alanine we applied (0.5 mM) significantly increased proliferation, whereas higher concentrations (2.5 or 5 mM) or the addition of PEG-CAT had adverse effects (Figure 1F). Of note, the pro-proliferative of D-alanine at low concentrations and the anti-proliferative effect at higher concentrations highlights the dual role of H_2_O_2_, which strongly depends on its dose and the antioxidant capacity of the experimental system (Gough & Cotter 2011).

**Figure 1.**
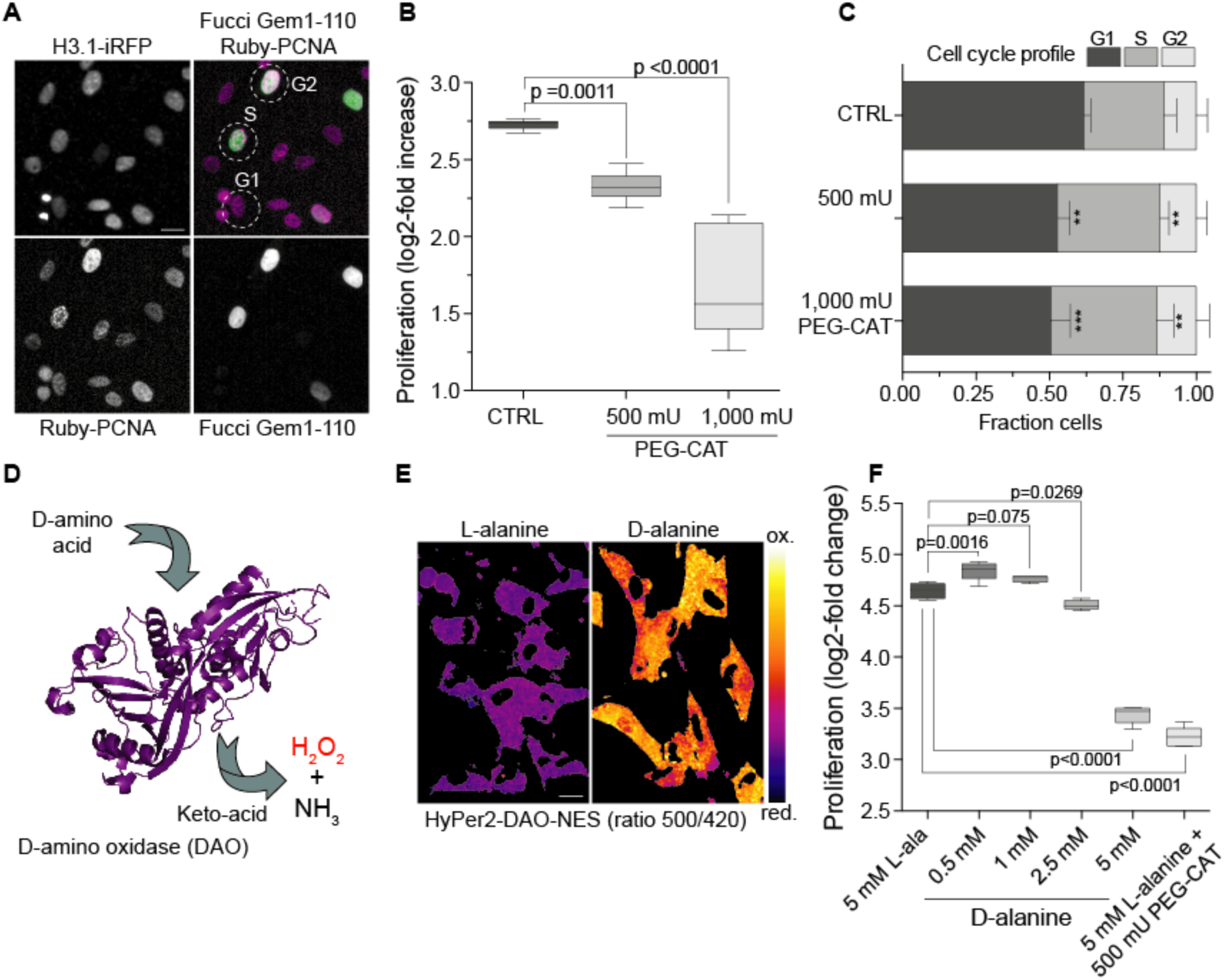
Proliferation and S phase progression require oxidative events in RPE-1. (A) Exemplary cell cycle analyses of single cells using endogenously tagged H3.1-iRFP for segmentation and Fucci-Gem1-110 and Ruby-PCNA to unambiguously classify cells, respectively. Fucci-Gem1-110 negative cells are in G1 phase, Fucci-Gem1-110 positive cells are in S or G2 phase, and PCNA-foci identify S phase. Examples of cell cycle classifications are indicated in the overlay image. Scale bar = 10 µm. (B) Proliferation of RPE-1 cells in the presence of the indicated units of PEG-catalase (PEG-CAT). Boxplots indicate the median log2-fold increase of cells for 48 hours PEG-CAT treatment. Significance according to one-way ANOVA with Dunnett’s multicomparison test (n=3, N=9). (C) Stacked bars showing the mean ± SD cell cycle phase distribution of cells from (B) and classified as in (A). Significance according to one-way ANOVA with Holm-Sidak’s multicomparison test: **(500 mU PEG-CAT(G1), p = 0.0011), ***(500 mU PEG-CAT(S), p = 0.0032), ***(1,000 mU PEG-CAT(G1), p = 0.0001), ****(1,000 mU PEG-CAT(S), p = 0.0012), (n=3, N=9). (D) Illustration of D-amino acid oxidase (DAO)-mediated H_2_O_2_ production in response to D-alanine. Structure from PDB:1COI without ligands. (E) Representative images showing ratio imaging of RPE-1 cells stably expressing cytoplasmic HyPer2-DAO-NES one hour after the addition of 10 mM D- or L-alanine. Scale bar = 10 µm. (F) Proliferation of RPE-1 cells expressing HyPer2-DAO-NES in the presence of the indicated concentrations of D- and L-alanine or L-alanine and PEG-CAT together. Boxplots indicate the median log2-fold increase of cells during 48 hours of treatment. Significance according to one-way ANOVA with Dunnett’s multicomparison test (N=5, data are representative of 3 independent experiments).

We conclude that oxidative events, mediated in particular by H_2_O_2_, are required for normal proliferation in RPE-1 cells. ROS appear to have a role in S phase progression indicating that non-transformed RPE-1 cells are ideally suited to reveal the molecular interplay between cell cycle progression and ROS signaling.

### ROS levels oscillate during the cell cycle

As ROS are required for proliferation in RPE-1 cells, we investigated how physiological ROS production is correlated with cell cycle progression. To this end we employed the ROS probe CellRox Deep Red to determine the overall ROS content in living cells expressing Ruby-PCNA and cyclin A2-Venus to simultaneously define cell cycle phases. We observed that compared to G1 phase, cells in S and G2 phases displayed increased labeling of CellRox Deep Red (Figure 2A). Quantification of thousands of cells by flow cytometry analysis revealed significant increases of cellular ROS from G1 to S and S to G2/M phases indicative of ROS oscillations during the cell cycle (Figure 2B). Intriguingly, the ∼25% increase of ROS from G1 to S phase is of the same magnitude as pro-proliferative ROS production by HyPer2-DAO-NES stimulated with 0,5 mM D-alanine (∼20%, Figure S1E). Our measurements were performed at atmospheric oxygen concentration (21%), which might affect the redox status of cells that experience lower oxygen concentration in their in vivo tissue context. Thus, we repeated our measurements using RPE-1 cells grown at an oxygen concentration reported to be typical for the retinal epithelium (6.3%). This confirmed our previous findings and indicated an even steeper increase of ROS from G1 to S phase (Figure 2C). Measuring ROS in primary human foreskin fibroblasts (BJ) grown at their reported *in vivo* oxygen concentration of 4% gave the same result (Figure 2D)(Balin et al. 2002).

**Figure 2.**
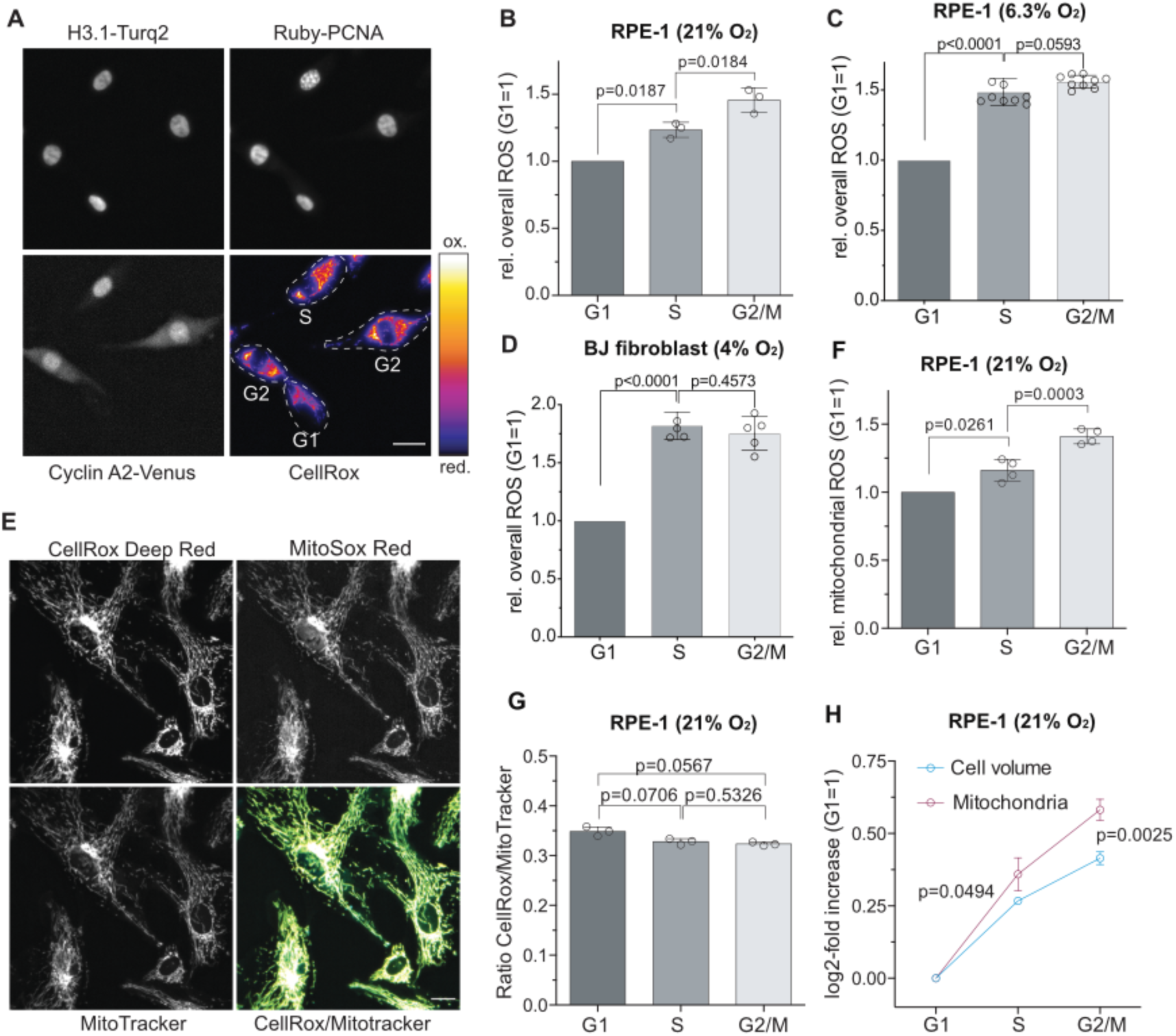
Mitochondrial ROS increases in S and G2 phase. (A) Representative images of living RPE-1 cells lablled with the ROS dye CellRox Deep Red illustrate endogenous overall ROS production during the cell cycle. Cell cycle classification based on endogenous cyclin A2-Venus and Ruby-PCNA (see also Figure S1B) are indicated. Scale bar = 25 μm. (B and C) Flow cytometry analysis of RPE-1 cells grown in normoxia (21% O_2_) or hypoxia (6.3% O_2_) labeled with CellRox Deep Red and Hoechst 33342. Bars represent the mean ± SD of the relative increase in CellRox Deep Red intensities normalized to G1 phase. Significances according to one-sample t-test (G1 versus S) and a two-tailed paired t-test (S versus G2) (B, n=3, N=3; C, n=3, N=8). (D) Analogous analyses as in (C) but with primary foreskin BJ fibroblast grown at 4% O_2_. (E) Representative images of living RPE-1 cells co-labeled with CellRox Deep Red, MitoSox Red and MitoTracker Green to quantify all mitochondria. Overlay indicate CellRox (yellow) and MitoTracker (cyan) co-localization. Scale bar = 25 μm. (F) Flow cytometry analysis of living RPE-1 cells stained with MitoSox Red and Hoechst 33342. Bars represent the mean ± SD of the relative increase in MitoSox Red intensities normalized to G1 phase. Significances according to one-sample t-test (G1 versus S) and a two-tailed paired t-test (S versus G2) (n=4, N=4). (G) Flow cytometry analysis of living RPE-1 cells co-labelled with CellRox Deep Red, MitoTracker Green and Hoechst 33342. Bars represent the mean ± SD CellRox/MitoTracker ratio. Significance according to paired one-way ANOVA with Holm-Sidak’s multi comparison test (n=3, N=3). (H) Quantification of the cell cycle increase in cell size based on forward scatter and mitochondria based on MitoTracker Green labelling. Data indicate the mean log2-fold increase ± SD of cell volume and mitochondrial mass, normalized to G1 phase. Significances according to multiple t-tests with Holm-Sidak’s correction for multiple comparisons (n=3, N=3).

Notably, the topology of CellRox Deep Red labeling appeared to be reminiscent of mitochondrial networks. Co-labelling of living cells with CellRox Deep Red and the mitochondrial marker MitoTracker Green displayed overlapping signals (Figure 2E) suggesting that the cell cycle-correlated ROS dynamics we detected likely recapitulate changes of mitochondrial ROS. Indeed, specific detection of mitochondrial ROS with the dye MitoSox Red mirrored localization and cell cycle-correlated dynamics of overall ROS (Figures 2E and 2F). The increase of mitochondrial ROS in S and G2/M could reflect an increase of mitochondrial activity, i.e. due to a metabolic switch from glycolysis to oxidative phosphorylation, and/or an increase of mitochondria mass in relation to the cell volume. To distinguish between both possibilities, we performed flow cytometry analysis labeling the overall mitochondrial content of living cells with the membrane-potential independent dye MitoTracker Green and mitochondrial ROS with MitoSox Red. An equal ratio of MitoSox Red to MitoTracker Green in each cell cycle phase implied that the increase of mitochondrial ROS predominately reflected an increase in the number of mitochondria (Figure 2G). As others before (Havens et al. 2006), we observed that the amount of mitochondria increased faster than the cell volume (Figure 2H) providing one explanation for the elevated concentrations of cellular ROS in S and G2 or M phases.

Taken together, our data reveal periodical oscillations of overall ROS during the cell cycle in both, normoxic and hypoxic tissue cell culture conditions that correlate with mitochondrial ROS dynamics. Our data imply that the increase in cellular ROS largely results from a relative increase in the number of mitochondria in S and G2/M compared to G1 phase.

### Mitochondrial ROS drives progression through S phase

Mitochondrial ROS are produced as a by-product of oxidative phosphorylation. Thus, reducing the influx of metabolites into the TCA cycle should decrease the amount of O2^•−^ released from the ETC, and consequently the concentration of mitochondria-derived H_2_O_2_ in the cytoplasm. In line with this idea, we employed endoribonuclease-prepared siRNAs (esi-RNAs) to downregulate PDHB, a subunit of the mitochondrial pyruvate dehydrogenase (PDH) complex that converts pyruvate to acetyl-CoA and acts as a gatekeeper between glycolysis in the cytosol and oxidative phosphorylation in mitochondria (Figure 3A). Indeed, depleting PDHB protein levels to below 25% (Figures 3B and 3C) was sufficient to significantly reduce cellular H_2_O_2_ detected by HyPer7, a ratio-metric H_2_O_2_ sensor with improved sensitivity (Pak et al. 2020) (Figure 3D). Importantly, reducing ROS production via PDHB depletion did not affect the ADP/ATP ratio indicating that cells were able to maintain ATP levels likely by increasing the glycolytic flux (Figure 3E).

**Figure 3.**
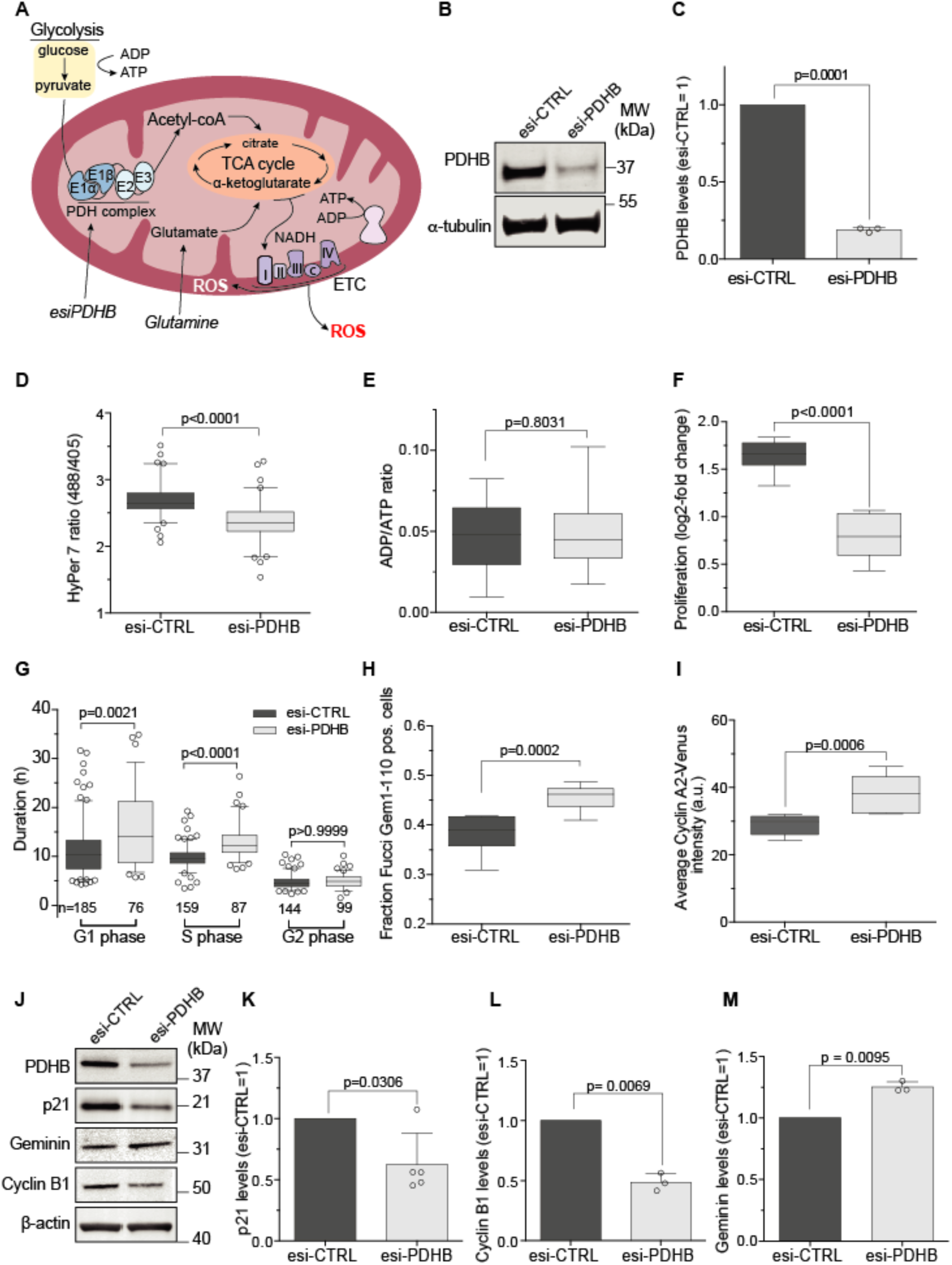
Mitochondrial ROS drive progression through S phase. (A) Illustration of mitochondrial metabolism showing metabolite and ROS flux and the PDH complex as the gatekeeper between glycolysis and the TCA cycle. (B) Representative Western blot analysis of cells treated for 48 hours with control (esi-CTRL) and esiRNAs targeting the β-subunit of the PDH complex (esi-PDHB). (C) Quantification of the degree of PDHB depletion from experiments in (B) normalized to esi-CTRL. Bars indicate the mean ± SD. Significance according to two-tailed one-sample t-test (n=3, N=3). (D) Analysis of endogenous ROS in RPE-1 cells stably expressing the H_2_O_2_ sensor HyPer7. Boxplots indicate the median HyPer7 ratio after treatment as in (B). Significance according to two-tailed unpaired t-test (n=3, esi-CTRL N=80, esi-PDHB N=82). (E) ATP levels quantification in response to PDHB depletion as in (B). Boxplots indicate the median ADP/ATP ratio. Significance according to two-tailed unpaired t-test with (n=3, N=12 esi-CTRL, N=11, esi-PDHB). (F) Proliferation analysis of PDHB depleted cells. Boxplots indicate the median log2-fold increase in the number of cells after treatment as in (B). Significance according to two-tailed unpaired t-test (n=3, N=9). (G) Cell cycle analysis showing duration of cell phases determined by single-cell time-lapse microscopy after treatment as in (B). Boxplots indicate the median duration cells require to progress through the indicated cell cycle phase. Significance according to Kruskal-Wallis with Dunnett’s multi comparisons test. (n=3, N is indicated in the graph). (H) Quantification of cells in S/G2 phase after treatment as in (B) Boxplots indicate the median fraction of Fucci-Gem1-110 positive cells. Significance according to two-tailed unpaired t-test (n=3, N=9). (I) Quantification of endogenous cyclin A2-Venus expression levels as a measure of S/G2 phase progression. Boxplots indicate the median intensity of cyclin A2-Venus after treatment as in (B). Significance according to two-tailed unpaired t-test (n=3, N=9). j, Representative Western blot analysis of cells treated as in (B) detecting cell cycle markers. (K-M) Quantification from data as in (J). Bars represent the mean ± SD normalized to esi-CTRL. Significance according to two-tailed one-sample t-test (n=3, N=5 (p21), N=3 (Cyclin B1, Geminin).

PDHB-depleted cells proliferated slower (Figure 3F) but did not induce apoptosis as judged by PARP cleavage (Figure S1F). In support of our results from reductive treatments with NAC and PEG-catalase (Figure 1) PDHB-depleted cells required significantly longer to progress through S phase in single-cell time-lapse microscopy (Figure 3G). We noticed that PDHB depletion in this experimental setup caused a prolonged and more heterogeneous duration of G1 phase. Unchanged levels of ATP and decreased levels of p21 (see below), however, indicated that longer G1 phases were not caused by activation of a metabolic checkpoint that arrests cells at the G1/S transition (Jones et al. 2005; Mitra et al. 2009). An alternative possibility explaining this result is technical as the computational cell cycle analyses pipeline employs PCNA foci to precisely stage the onset of S phase (Zerjatke et al. 2017). Since PDHB depletion reduces the number of PCNA foci in S phase (see below and Figure 4) our analysis likely under-estimated the duration S phase in favor of G1 phase. As orthogonal approaches, we assessed the cell cycle distribution of PDHB depleted cells by Western analyses for established cell cycle markers or by single-cell analyses for the S phase markers cyclin A2 and Fucci Gem1-110. Quantitative Western blot analysis of total extracts (Figure 3J) showed that the protein levels of G1 and G2 phase markers p21 and cyclin B1 were lower in PDHB depleted cells (Figures 3K and 3L), whereas the S phase protein geminin accumulated (Figure 3M). In agreement with a delay in S phase but not in G1 phase, PDHB depletion increased the proportion of Fucci Gem1-110 positive cells (Figure 3H) as well as the intensity of cyclin A2-Venus (Figure 3I), two markers that only accumulate after the G1/S transition.

**Figure 4.**
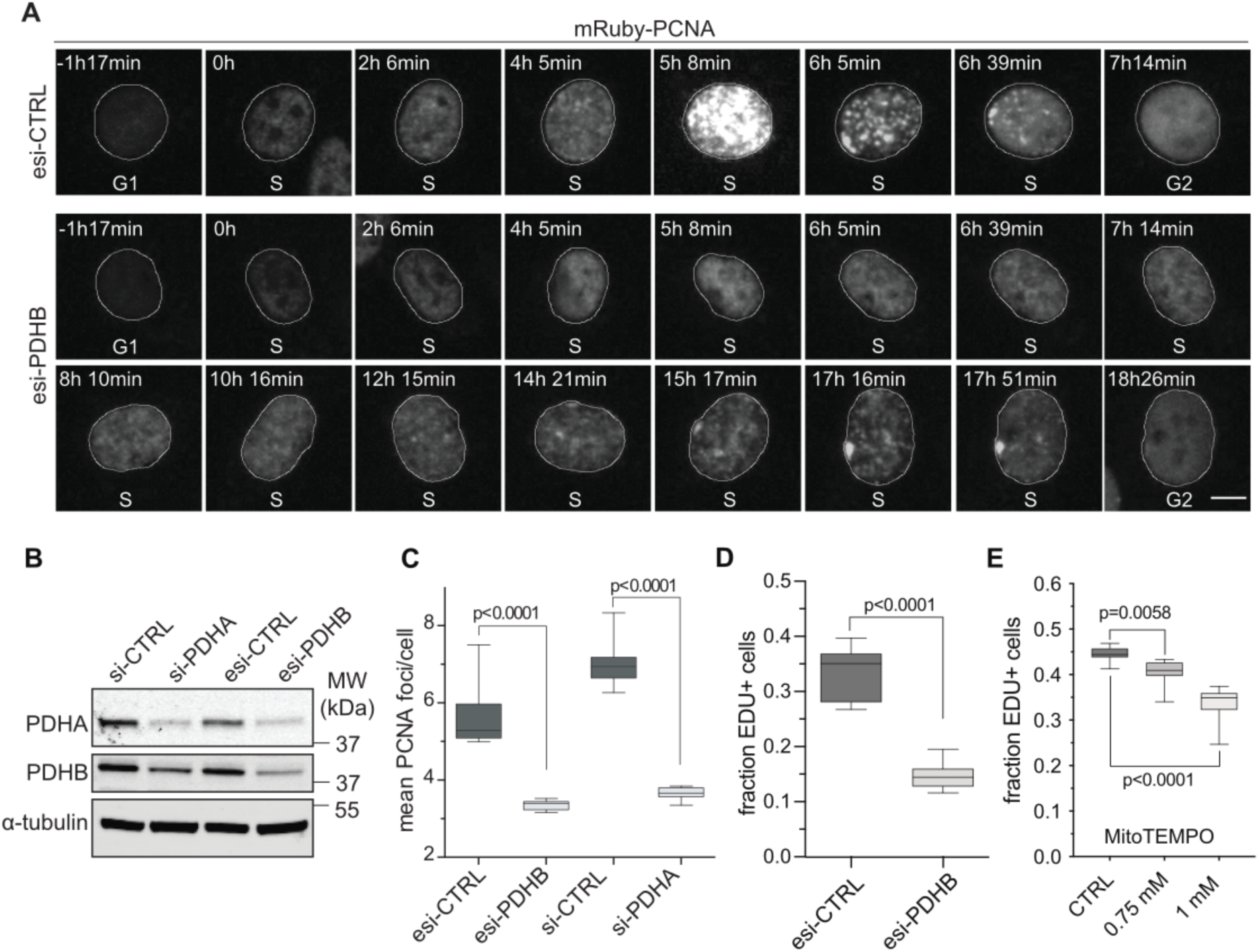
Mitochondrial ROS promote DNA replication. (A) Montage of time-lapse images from the beginning to the end of S phase (T=0) highlighting PCNA foci formation in control (esi-CTRL) and PDHB-depleted cells (esi-PDHB) between 24 and 72 hours after treatment. Scale bar = 20 µm. (B) Representative Western blot analysis of cells depleted for 48 hours of PDHA (si-PDHA), PDHB (esi-PDHB) or controls (si-CTRL, esi-CTRL). Note, targeting one PDH complex subunit results in partial co-depletion of the other. (C) Quantification of PCNA foci in S phase in cells treated as in (B). Boxplots indicate the median number of detectable PCNA foci. Significance according to two-tailed unpaired t-test (n=3, N=9). (D) Cells treated as in (A) were exposed for 45 minutes to 10 µM EDU to detect DNA replication and analyzed by single-cell imaging. Boxplots indicate the median fraction of EDU-positive cells. Significance according to two-tailed unpaired t-test (n=3, N=18). (E) EDU incorporation as in (D) but with RPE-1 cells treated for 48 hours with the indicated concentrations of MitoTEMPO. Boxplots indicate the median fraction of EDU-positive cells. Significance according to two-tailed unpaired t-test (n=4, N=15: CTRL, 1mM, n=2, N=9: 0,75 mM).

We conclude that decreasing the metabolic flux through the TCA cycle via interference with the PDH complex reduces the concentration of cellular H_2_O_2_ resulting in slower cell proliferation. The increased time required to progress through S phase and the accumulation of S, but not of G1 or G2 phase markers suggests that a major cell cycle-associated role of mitochondrial ROS lies in S phase.

### Mitochondrial ROS promotes DNA replication

At the beginning of S phase replication machines cluster into replication factories corresponding to microscopically-visible PCNA foci (Leonhardt et al. 2000). Since reducing mitochondrial ROS levels via PDHB depletion slowed S phase, we monitored the intensity and number of PCNA foci during time-lapse imaging as a surrogate to assess the fidelity of DNA replication. Replication foci in control cells followed the stereotypical spatiotemporal pattern described for replication factories (Leonhardt et al. 2000): multiple small foci in early S phase, concentration of foci in mid S phase, and formation of large clusters at the nuclear periphery in late S phase (Figure 4A and Movie 1). In contrast, PCNA foci were hardly detectable until the very end of S phase in PDHB depleted cells at the magnification used (Figure 4A and Movie 2). To exclude potential off-target effects of esi-PDHB treatment, we targeted another subunit of the PDH complex, PDHA (Figure 4B). Quantifying the mean number of PCNA foci in S phase cells confirmed that depleting either PDHA or PDHB strongly decreased the formation of detectable PCNA foci (Figure 4C). To strengthen the link of mitochondrial ROS and DNA replication we also assessed EDU incorporation in PDHB depleted cells or cells treated for 48 hours with the mitochondrial antioxidant MitoTEMPO. Indeed, both treatments significantly reduced the fraction of EDU positive cells compared to control (Figures 4D, E)

Taken together these data show that metabolic interference with the TCA cycle or mitochondrial antioxidant treatment affect replication factory formation and DNA replication during S phase. Since interference with the PDH complex reduces cellular H_2_O_2_ and hinders DNA replication, we conclude that mitochondrial ROS promote DNA replication.

### CDK2 activity and T-loop phosphorylation are sensitive to mitochondrial ROS

CDK2 mediates not only the initiation of DNA replication but also regulates the spatiotemporal pattern of replication in the nucleus (Sansam et al. 2015). Since we observed changes in PCNA foci formation and dynamics in response to PDH complex perturbation (Figure 4), we investigated whether mitochondrial ROS promotes CDK2 activity in S phase. To assess CDK2 activity in living cells, we created a RPE-1 cell line stably expressing a CDK2 activity sensor that localizes to the nucleus upon low CKD2 activity, and becomes progressively more cytoplasmic once CDK2 activity increases (Spencer et al. 2013) (Figure 5A). Depleting PDHB for 48 hours strongly increased the number of cells with nucleoplasmic localized CDK2 sensor (Figure 5B and 5C) and thus the percentage of cells with low CDK2 activity (CDK2^low^) from 34% to 61%. Note that CDK2^low^ cells are unlikely to be in quiescence or arrested before the restriction point (Spencer et al. 2013) because the same treatment resulted in accumulation of multiple cell cycle markers that are expressed downstream of the G1/S transition, including cyclin A2 and geminin (Figure 3). Next, we addressed how mitochondrial metabolism and ROS production affect CDK2 activity on the molecular level. To reach full activity CDK2 requires binding by E or A-type cyclins and phosphorylation of T160 within its T-loop. Because PDHB depleted cells can enter S phase (increased proportion of Fucci-Gem1-110 positive cells, Figure 3H) and express cyclin A2 (Figure 3I), we investigated whether T160 phosphorylation was affected. Indeed, quantitative Western blot analysis of lysates from control and PDHB- depleted cells with a phosphorylation-specific T160 antibody (CDK2-pT160) showed a ∼50% reduction in T-loop phosphorylation (Figure 5D and 5E). Importantly, this difference was not due to an enrichment of PDHB-depleted cells in G1 phase since repeating the experiment in cells synchronized at the beginning of S phase with thymidine gave a comparable result (Figure S2A and S2B). To confirm these data by an alternative approach, we starved cells for 6 hours of glutamine, which is an established approach to reduce mitochondrial ROS production (Oh et al. 2020). As observed for depleting PDHB, 6 hours of glutamine starvation strongly increased the number of CDK2^low^ cells in both, normoxic (21% O_2_) and hypoxic (6.3% O_2_) culture conditions (Figure 5F, I) and reduced T160 phosphorylation (Figures 5G, H and 5J, K). Importantly, 6 hours of glutamine starvation did not synchronize cells in G1 phase, where pT160 is lower (Figure S2C). Finally, the addition of 8 mM and 10 mM NAC for 5 hours to S phase cells also reduced T160 phosphorylation, though to a lesser extent (Figures S2D and S2E). Exposure of cells to different concentrations of NAC for 48 hours as in proliferation experiments in Figure S1 decreased pT160 by up to 83% (Figure S2G) and resulted in CDK2 sensor accumulation in nucleus indicative of reduced CDK2 activity (Figure S2F). Importantly, the fraction of cyclin A2 positive cells increased upon NAC treatments in agreement with a delay in S but not in G1 phase (Figure S2H).

**Figure 5.**
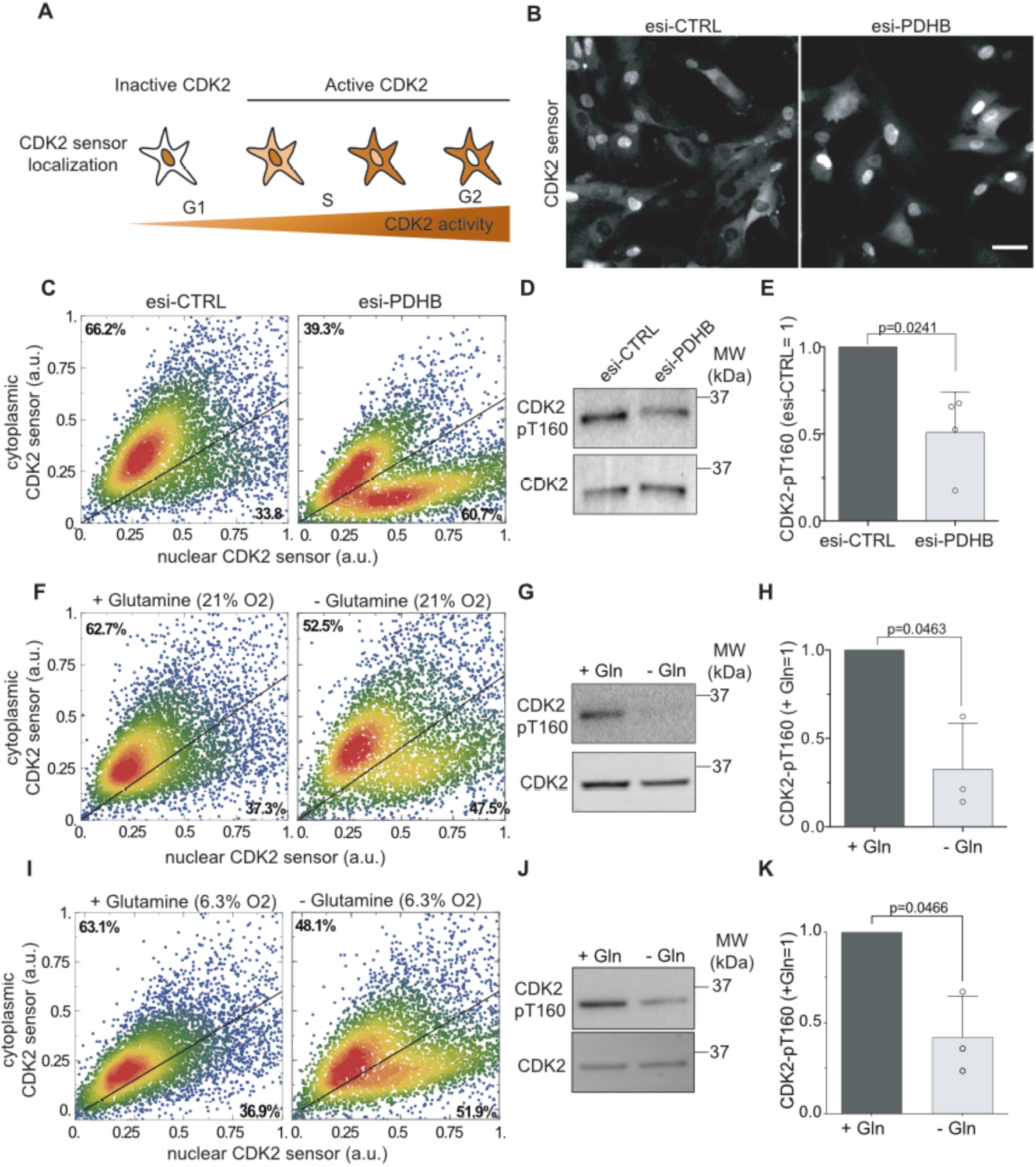
CDK2 activity and T-loop phosphorylation is sensitive to mitochondrial ROS. (A) Illustration showing the CDK2 activity-dependent localization of the CDK2 sensor. (B) Representative images of RPE-1 cells stably expressing the CDK2 sensor 48 hours after esi-CTRL and esi-PDHB transfection. Note the contrast of each image was adjusted separately to prevent saturation of nuclear -localized CDK2 sensor in esi-PDHB treatment. Scale bar = 50 µm. (C) Scatter plots show scaled intensities of cytoplasmic and nucleoplasmic localized CDK2 sensor of single cells treated as in (B). The percentage of CDK2^Low^ and CDK2^High^ is indicated in the plot (n=2, N=6,000). (D) Representative Western blot analysis detecting the degree of T160 phosphorylation of cells treated as in (B). (E) Quantification of the data shown in (D). Bars indicate the mean ± SD. Significance according to two-tailed one-sample t-test (n=4, N=4). (F) Scatter plots show scaled intensities of cytoplasmic and nucleoplasmic localized CDK2 sensor of single cells with or without glutamine (Gln) after 6 hours treatment grown in normoxia (21% O_2_) (n=3, N=6,000). (G) Representative Western blot analysis detecting the degree of T160 phosphorylation of cells treated as in (F). (H) Quantification of the data shown in (G). Bars indicate the mean ± SD. Significance according to two-tailed one-sample t-test (n=3, N=3). (I) Analysis as in (F) but with RPE-1 cells grown at 6.3% O_2_ (n=3, N=6,000). (J-K) Representative Western blot analysis and quantification of T160 phosphorylation from cells grown in 6.3% O_2_ as in (G-H). Bars indicate the mean ± SD. Significance according to two-tailed one-sample t-test (n=3, N=3).

Taken together our data show that reducing conditions achieved either by metabolic perturbation of oxidative phosphorylation or by a chemical reductant, decreased T-loop phosphorylation of CDK2. Hence, the degree of CDK2 activity could be coupled to mitochondrial metabolism using ROS as a messenger.

### Genetically-enabled ROS production increases T-loop phosphorylation

Thus far, our experiments in reducing conditions demonstrated indirectly that ROS was involved in regulating CDK2 activity via T160 phosphorylation. To provide more direct evidence, we tested whether we could rescue the loss of T160 phosphorylation after PDHB depletion by DAO-mediated H_2_O_2_ production (Figure 1). As CDK2-cyclin E/A complexes are predominately nucleoplasmic, we created a RPE-1 cell line stably expressing DAO fused to a nuclear localization sequence (NLS, HyPer2-DAO–NLS) to rapidly generate ROS in response to addition of D- but not to L-alanine (Figures 6A and 6B). Based on the gradual increase in the HyPer2 ratio (Figure 6B), we induced H_2_O_2_ for one and two hours and analyzed the consequences for T160 phosphorylation in PDHB-depleted cells. One hour of nucleoplasmic H_2_O_2_ production was sufficient to increase T160 phosphorylation significantly (Figures 6C and 6D). Because mitochondria-derived H_2_O_2_ needs first to pass the nuclear membrane to reach CDK2-cyclin E/A complexes, we repeated the experiment with cytoplasmic HyPer2-DAO–NES (Figure 1H). We found that cytoplasmic H_2_O_2_ production also rescues T160 phosphorylation (Figures 6E and 6F) indicating that indeed mitochondrial ROS can target predominantly nuclear-localized CDK2.

**Figure 6.**
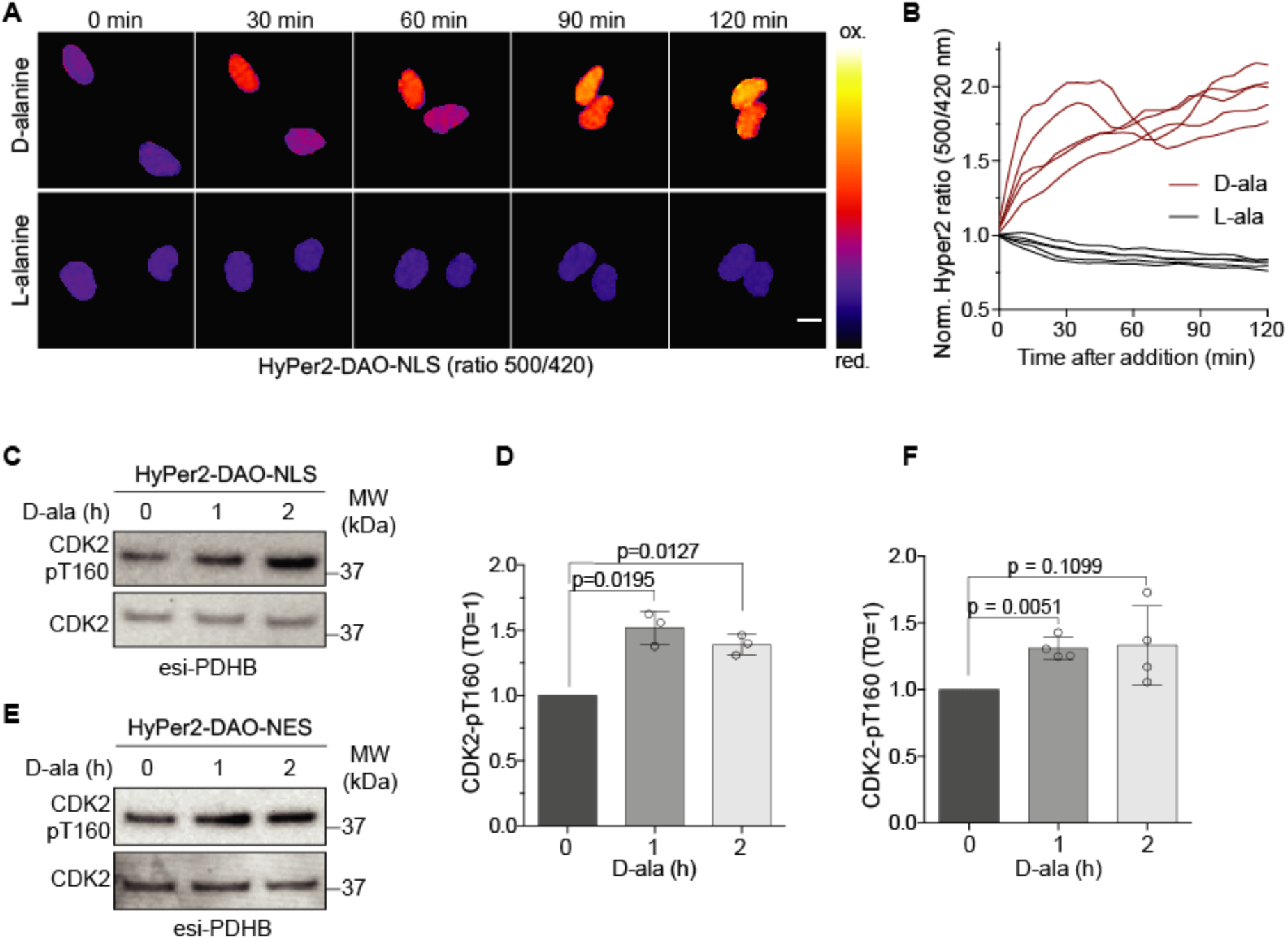
Genetically-enabled ROS production increases T-loop phosphorylation. (A) Representative time-lapse images showing ratio imaging of RPE-1 cells stably expressing nucleoplasmic localized HyPer2-DAO-NLS during 2 hours of treatment with 10 mM D- or L-alanine (D-ala, L-ala). Scale bar = 10 µm. (B) ROS production of HyPer2-DAO-NLS in response to D-ala and L- ala addition. Data show 5 independent cell tracks per condition normalized to ratio at the time of D-ala or L-ala addition (T0=1). (C) Representative Western blot analysis showing an increase of T160 phosphorylation in PDHB-depleted cells after D-ala (10 mM)-induced production of ROS by HyPer2-DAO-NLS. (D) Quantification of data shown in (C). Bars represent the mean ± SD normalized to the time of D-ala addition (T=0). Significance according to two-tailed one-sample t-test. (n=3, N=3). (E) Representative Western blot analysis as in (C) but with cytoplasmic ROS production by HyPer2-DAO-NES. (F) Quantification data as in (E). Bars represent the mean ± SD normalized to time of D-ala addition (T=0). Significance according to two-tailed one-sample t-test. (n=4, N=4).

We conclude that genetically-enabled H_2_O_2_ production promotes T-loop phosphorylation on CDK2. The observation that cytoplasmic production of H_2_O_2_ can target nucleoplasmic CDK2 supports the idea that mitochondrial ROS acts as a regulator of CDK2 activity.

### Preventing C177 oxidation increases KAP binding to CDK2

Our finding that ROS stimulates CDK2 T-loop phosphorylation led us to hypothesize that either CDK2 itself or its T160 phosphorylation regulating enzymes, CAK and KAP, are targeted by ROS. To evaluate these possibilities, we took advantage of the membrane-permeant chemo-selective probe BTD to label sulfenic acids in living cells (Gupta & Carroll 2016). We added BTD for 30 minutes to RPE-1 cells, followed by fixation in paraformaldehyde (PFA) and simultaneous quenching of all unreacted cysteines with iodoacetamide (IAA). Subsequently, we permeabilized cells, clicked azide-biotin to the alkyne moiety of BTD, reversed the PFA crosslink by boiling, and enriched for sulfenic acid-modified proteins by streptavidin pull downs (Figures 7A and S2J). In Western blots from denaturing streptavidin pull downs, we detected CDK2 and the known sulfenic acid-modified protein GAPDH but not the highly abundant ribosomal protein RPL26, which does not contain cysteine residues (Figure 7B). In agreement, several mass spectrometry studies using BTD or alternative probes in H_2_O_2_-treated cell extracts or unperturbed cells identified CDK2 oxidation to sulfenic acid only on C177 (Yang et al. 2015; Gupta et al. 2017; Xiao et al. 2020; Shi et al. 2021). CDK2 contains three cysteine residues but the crystal structure of CDK2 indicates that only C177 is solvent-exposed (Brown et al. 1999) and thus can be a target of oxidation. Even though, C177 is the only cysteine identified by proteomic studies, we cannot exclude that the other two cysteines of CDK2 can be oxidized as well. C177 is positioned in an unstructured loop in close proximity to T160 (Figure S2K), is conserved in vertebrates (Figure S2L), and is only found in CDK2 and not in other related human CDKs (Figure 7C). Next, we assessed whether CDK2 oxidation increased from G1 to S and G2 phase as predicted by the increasing ROS levels during the cell cycle. We expressed CDK2 tagged at its C terminus with the influenza hemagglutinin epitope (HA) in cells synchronized in G1, S and G2 by a release from 24 hours of serum starvation for 8, 18 and 22 hours, respectively (Figure S2I). After 30 minutes of labelling of living RPE-1 cells with BTD, we lysed cells under reducing conditions followed by clicking the fluorescent dye Az800 to BTD (Figure S3A, B). Subsequent detection of Az800 on CDK2-HA immuno-precipitates revealed that cysteine oxidation on CDK2 to sulfenic acid increased from G1 phase to S and G2 phase in a manner reflecting the overall increase of cell cycle ROS (compare Figure 7D and Figure 2).

**Figure 7.**
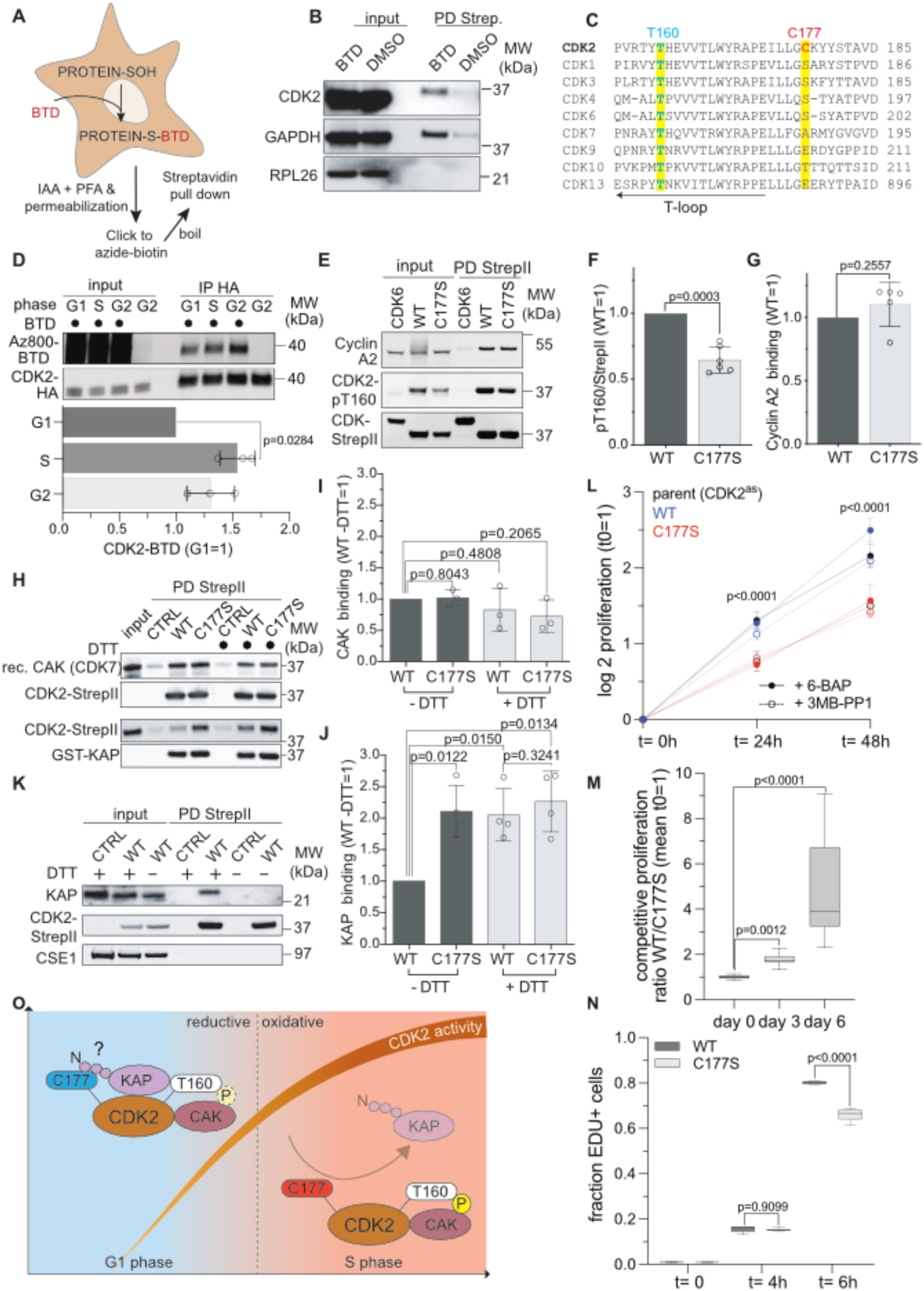
Preventing C177 oxidation increases KAP binding to CDK2. (A) Illustration showing the purification of sulfenic acid modified-proteins (Protein-SOH) in living cells with the chemo-selective probe BTD. Experimental details are described in the main text and methods. (B) Representative Western blot analysis (n=3) of denaturing Streptavidin pull downs with CDK2, GAPDH (positive control) and RPL26 (negative control) antibodies from cell lysates prepared from control (DMSO) and BTD-treated RPE-1 cells as illustrated in (A). Note, the corresponding detection for biotin is presented in Figure S3A. (C) Clustal Omega alignment of the T-loop and downstream sequence of CDKs in *H. sapiens*. Residues aligning to T160 (blue) and C177 (red) of CDK2 are highlighted in yellow. (D) Representative scan and quantification of an SDS-PAGE gel detecting sulfenic acids on immuno-purified CDK-HA from BTD lablled cells synchronized into G1, S and G2 phases. BTD detection was performed by click chemistry with the fluorescent dye Az800 and normalized to the amount of CDK2-HA. Significance according to two-tailed one-sample t-test. Scans showing the complete gel at lower intensities to judge the inputs are presented in Figure S3B (n=3, N=3). (E) Representative Western blot analysis of StrepII pull downs using cell extracts from S-phase synchronized RPE-1 cells transiently expressing CDK2-WT-StrepII (WT), CDK2-C177S-StrepII (C177S) or CDK6-StrepII as a control. The relative degree of T160 phosphorylation (pT160) and co-precipitated cyclin A2 normalized to WT is shown beneath the blot. (F-G) Quantification of pT160 (F) and cyclin A2 binding (G) from data shown as in (E). Bars represent the mean ± SD. Significance according to two-tailed one-sample t-test. (pT160: n=6, N=6; Cyclin A2: n=4, N=4). (H) Representative Western blot analysis of an *in vitro* binding assay between recombinant KAP or CAK and CDK2-StrepII purified from S-phase cells. The relative binding of recombinant CAK or KAP to CDK2-StrepII in non-reducing (-DTT) and reducing conditions (+10 mM DTT) normalized to WT (-DTT) are shown below the blot, respectively. A silver analysis of Strep II pulldown used for the experiments is shown in Figure S3H. Note, for CAK binding assays CDK2 was dephosphorylated with λ-phosphatase (see Figure S3I) to facilitate CDK7 binding. (I-J) Quantification of CAK or KAP binding to CDK2-StrepII from data as in (H). Bars represent the mean ± SD. Significance according to two-tailed unpaired one-sample t-test. (CAK: n=3, N=3; KAP: n=4 N=4). (K) Representative Western blot analysis showing the binding of endogenous KAP to CDK2-StrepII purified from S-phase cells. Note, before extract preparation (+DTT) cells were treated for 10 min with 5 mM DTT and the pull down was performed in the absence (-DTT) or presence of 20 mM DTT (+DTT) (n=3). (L) Representative proliferation assay (n=3) of RPE-1 CDK2^as^ cells expressing ectopic CDK2 WT or C177S in the presence of 10 µM inhibitory ATP analog 3MB-PP1 or 0.5 µM control 6-BAP. Note, adding 3MB-PP1 inactivates endogenous CDK2^as^ but not ectopic CDK2. Significance according to two-way ANOVA with Dunnett’s correction for multiple comparison. For clarity, only the values comparing 3MB-PP1 treated WT and C177S cells are indicated (N=12). (M) Competitive growth analyses of CDK2 WT and C177S cells in the presence of 3MB-PP1 seeded into the same well. WT and C177S were identified at the indicated time points by image analysis detecting GFP or mRuby expressed from the same mRNA as CDK2, respectively. Boxplots indicate the median ratio of GFP/mRuby cells normalized to the mean at the beginning of the experiment (t=0). Significance according to one-way ANOVA with Dunnett’s correction for multiple comparison (n=3, N=18). (N) Representative (n=3) EDU incorporation assay (10 µM) with CDK2 WT and C177S cells in the presence of 3MB-PP1 at the indicated time after release from a 24-hour G1 phase block with 150 nM Palbociclib. Note, EDU was added to t=0 cells during the last hour of Palbociclib treatment. Boxplots indicate the median ratio fraction of EDU-positive cells. Significance according to one-way ANOVA with Sidak’s correction for multiple comparisons (N=6). (K) Model of ROS-regulated CDK2-KAP interactions during the cell cycle. In more reductive conditions in G1 phase, both KAP and CAK interact with CDK2 and can act on T160. Here, the N-terminus of KAP might recognize reduced C177 (blue). Oxidation of C177 (red) in response to increased mitochondrial ROS in S phase prevents KAP binding to CDK2 and shifts the balance towards CAK binding. This ensures full T160 phosphorylation and CDK2 activity even in the presence of KAP.

To investigate the role of C177 oxidation in T160 phosphorylation, we mutated C177 to cysteine-mimetic serine (C177S) or alternatively to alanine (C177A). We transiently transfected StrepII-tagged versions of these constructs into RPE-1 cells synchronized at the beginning of S phase. StrepII pull downs showed that phosphorylation of T160 on C177S (Figure 7E and 7F) and C177A (Figures S3C and S3D) mutants were strongly reduced, whereas the binding of cyclin A2 (Figures 7G and S3E) and cyclin E1 (Figures S3F and S3G) was not significantly affected.

A possible explanation for this result is that C177 oxidation regulates the binding of either CAK or KAP to CDK2 and thereby, the phosphorylation status of T160. To test this idea, we performed binding assays comparing the binding of recombinant CAK and KAP to CDK2-StrepII WT and C177S purified from S phase-synchronized cells, the time when CDK2 should be oxidized (Figure S3H). For CAK binding assays, we first dephosphorylated bead immobilized CDK2-StrepII WT and C177S using λ-phosphatase (Figure S3I) before adding recombinant CAK in the presence or absence of the reducing agent dithiothreitol (DTT) (Figure 7H). We did not observe significant changes in CAK binding to CDK2 as determined by quantitative Western blotting for CDK7 (Figure 7I). In contrast, when we assessed the binding of recombinant KAP to phosphorylated CDK2, we found that the C177S mutant bound about twice as much KAP as the WT (Figure 7H). Importantly, under reducing conditions (+ DTT) the binding of KAP to CDK2 WT but not to C177S increased (Figure 7J). Finally, we investigated whether endogenous KAP binds to CDK2 in a redox-dependent manner. To this end, we performed StrepII pull downs of CDK2 WT from S phase-synchronized cells treated without (- DTT) and with 5 mM DTT for 10 min (+DTT) before lysis. Indeed, endogenous KAP is only bound to CDK2 in reducing conditions but not when DTT was omitted before and during extract preparation (Figure 7K). In contrast, HA-tagged CDK2 C177S but not WT was able to precipitate endogenous KAP in the absence of DTT suggesting that mutating C177 is sufficient to render the CDK2-KAP interaction independent of the oxidation state of CDK2 (Figure S3J).

If CDK2 oxidation on C177 is important for full CDK2 activation and S phase progression as our data suggest then cells expressing C177S in the absence of endogenous CDK2 activity should be impaired in proliferation and DNA replication. As our results indicate that C177 is key to proliferation we were hesitant to directly introduce a C177S mutation into genomic CDK2. Instead, we created pools of 5 independent stable cell lines expressing untagged CDK2 WT or C177S in RPE-1 cells harboring two alleles of Shokat analogue sensitive CDK2 (CDK2^as^)(Merrick et al. 2008). This allowed us to rapidly inactivate endogenous CDK2 and assess whether C177S can promote proliferation and DNA replication comparably to WT CDK2. Indeed, the addition of 10 µM inhibitory Shokat ATP analog 3MB-PP1 but not of 0.5 µM non-inhibitory 6-benzylaminopurine (6-BAP) decreased the proliferation of parent CDK2^as^ cells as previously reported (Figure 7L) (Merrick et al. 2008). Cells expressing CDK2 WT were only mildly affected by 3MB-PP1 as expected since ectopic CDK2 is not analogue sensitive. In contrast, expression of C177S slowed proliferation to the extent of 3MB-PP1 treated parent CDK2^as^ cells, whether or not 3MB-PP1 was added. This suggest that the C177S mutant has dominant-negative properties as known from CDK T-loop mutants (van den Heuvel & Harlow 1993). In agreement, CDK2 WT cells proliferated faster in competition assays than C177S expressing cells (Figure 7M). This was not due to lless ectopic CDK2 because C177S was expressed slightly higher than WT (Figure S3K). Finally, we assessed the ability of C177S to promote DNA replication. To account for potential differences in cell cycle distribution of WT and C177S cells during asynchronous proliferation we synchronized cells in G1 phase by adding 150 nM of the CDK4/6 inhibitor palbociclib for 24 hours (Trotter & Hagan 2020). We released cells in the presence of EDU and probed for DNA replication 4 and 6 hours later, reflecting early and mid S phase, respectively (Figure S3M) (Trotter & Hagan 2020). In agreement with our conclusion that C177 oxidation promotes CDK2 activity and DNA replication C177S expressing cells incorporated significantly less EDU compared to WT (Figure 7N).

Taken together, these data demonstrate that oxidation of C177 positioned in close proximity to the T-loop is important to maintain full T160 phosphorylation and CDK2 activity. Mutation or reduction of C177 increases KAP but not CAK binding, indicating that ROS promote S phase progression and DNA replication by virtue of preventing T160 phosphatase recruitment.

## DISCUSSION

To proliferate, cells must coordinate cell growth driven by metabolism and cell cycle progression to ensure that DNA and other essential cellular components are duplicated before cell division. Principles of bidirectional cross-talk between cell cycle and metabolic regulation are just emerging and are an active area of research, not at least because aberrations in both systems are hallmarks of numerous diseases including cancer. In the presented study, we uncover that mitochondrial ROS continuously increase from G1 to S and G2 phases and oxidize CDK2. Oxidation occurs at a conserved cysteine residue in close proximity to the T-loop of CDK2 and prevents the binding of KAP. As a consequence, T-loop phosphorylation can be sustained despite high KAP expression during early S phase (Gyuris et al. 1993) guaranteeing full CDK2 activation and progression through S phase. Thus, we identify a simple but elegant mechanism by which cell cycle-dependent production of mitochondrial ROS promotes DNA replication to coordinate cell cycle progression with the metabolic state of the cell.

The concept that the redox environment of cells is coordinated with cell cycle progression dates back to 1931 when Rapkine (Rapkine 1931) postulated a thiol cycle in dividing urchin eggs. Since then, redox oscillations similar to the ROS dynamics we observe here have been inferred by several studies based on indirect experiments with reducing agents such as NAC (Menon et al. 2007; Kim et al. 2001; Kyaw et al. 2004; Havens et al. 2006) and directly using the ROS-sensitive fluorescent probe dichlorodihydrofluorescein diacetate DCFH-DA(Kim et al. 2001; Kyaw et al. 2004; Havens et al. 2006; Menon & Goswami 2007; Ivanova et al. 2021). However, ROS measurements with DCFH-DA are inherently prone to artifacts and have several limitations (Kalyanaraman et al. 2012; Bonini et al. 2006). Therefore, the source and characteristics of ROS oscillations during the cell cycle in proliferating cells have remained controversial. In our study, we have used non-transformed RPE-1 cells and the ROS probe CellRox Deep Red, which labels the hydroxyl radical, O2^•−^ and H_2_O_2_ in order of sensitivity. We find that CellRox Deep Red predominately labels mitochondrial structures in living cells, which is consistent with O2^•−^ production in mitochondria. Our data suggest that the cell cycle-dependent increase of cellular ROS likely recapitulates the relative increase in the number of mitochondria rather than increased respiration (Figure 2). We cannot, however, exclude that bursts of mitochondrial activity that are not detectable by our analyses also contribute to ROS production, e.g. during the G1/S transition (Mitra et al. 2009). Of note, the increase of ROS in S and G2 phase contradicts observations in yeast where DNA replication occurs in the reductive phase of the metabolic cycle (Tu et al. 2005; Chen et al. 2007). In animals and plants, however, both oxidative phosphorylation and glycolysis appear to be generally upregulated during the cell cycle to provide the metabolites and energy required for DNA synthesis and chromosome segregation (Salazar-Roa & Malumbres 2017). This difference may reflect that budding predominately reproduces as a haploid organism and thus requires a more protective reductive environment to avoid deleterious mutations during DNA synthesis.

Previous studies in *Drosophila* and human cells have suggested that mitochondria are needed to provide the energy and the buildup of cyclin E required to enter S phase (Mandal et al. 2005; Mitra et al. 2009; Robbins & Morrill 1969). Our work now identifies mitochondrial ROS as a key driver of S phase that ensures full CDK2 activity necessary for DNA replication (Figures 1, 3 and 4). How can these findings be reconciled and integrated into a congruent model of S phase regulation by mitochondria? We find no evidence that restricting the metabolic influx of acetyl-CoA into the TCA cycle decreases ATP levels and activates a metabolic checkpoint that restricts the entry in S phase in a p53 and p21-dependent manner (Figure 3) (Jones et al. 2005; Mitra et al. 2009). In agreement with this, mitochondrial ATP production appears to be dispensable in human cells with access to sufficient glucose (Sullivan et al. 2015). Towards the end of G1 phase mitochondria form a transient hyperfused network that, by an as yet to be defined mechanism possibly involving p53, boosts the levels of cyclin E to drive cells into S phase (Mitra et al. 2009). Our data indicate that the limiting function of ROS is downstream of this process because cells with decreased levels of mitochondrial ROS display elevated levels of the S and G2 phase markers geminin and cyclin A2 (Figure 3). Thus, mitochondria seem to employ two sequential and distinct mechanisms to regulate S phase via CDK2 activity: energy-driven expression of cyclin E to initiate S phase and ROS-sustained T-loop phosphorylation to drive DNA replication (see below).

Two studies have suggested that the critical function of mitochondria in proliferating cells is to synthesize aspartate as a building block for proteins and nucleotides (Sullivan et al. 2015; Birsoy et al. 2015). In this context, respiration performs an essential anabolic function and provides molecular oxygen as the terminal electron acceptor for aspartate synthesis during oxidative TCA metabolism. In respiration-incompetent cells with limited access to electron acceptors aspartate becomes limiting for nucleotide biosynthesis and cells cease proliferation in S phase (Sullivan et al. 2015). This raises the intriguing possibility that a major physiological function of mitochondrial ROS is to synchronize nucleotide and DNA synthesis by using CDK2 activity regulation as a speed dial of replication.

Even 40 years after the discovery of T-loop phosphorylation (Gould et al. 1991), a congruent model that can explain how CAK and KAP activities are coordinated to avoid futile cycles of phosphorylation and de-phosphorylation is lacking. In more complex organisms CDK2 and CDK1 are the only cell cycle regulatory CDKs that sequentially bind to different cyclins during the cell cycle and thus can become a target of KAP as monomers, i.e. during the degradation of cyclin E in early S phase and of cyclin A during prometaphase, respectively. However, in contrast to CDK2, CAK and KAP never compete for CDK1 because CDK1 is only phosphorylated by CAK once the cyclin is bound and in this state, the T-loop is inaccessible to KAP (Poon & Hunter 1995; Song et al. 2001). For CDK2, however, the T-loop becomes accessible to KAP once CDK2 switches from cyclin E to cyclin A at the beginning of S phase, and yet T160 phosphorylation remains constant until the end of mitosis (Figure S2I). Our finding that the interaction between CDK2 and KAP is regulated by oxidation of C177 (Figure 7) solves this issue and explains why T160 phosphorylation is sensitive to interference with mitochondrial ROS production (Figure 5). We propose a model by which the increase of mitochondrial ROS in S phase prevents KAP binding to CDK2 to ensure that T-loop phosphorylation of CDK2 can be sustained at times of high KAP expression (Gyuris et al. 1993) (Figure 7K). An exception to this rule is CDK2 in *Drosophila*; but here CDK2 does not switch cyclins because cyclin A only interacts with CDK1 (Harper & Elledge 1998). In support of this model, a cysteine in the position of C177 is absent in unicellular organisms and plants, where the function of CDK2 is carried out by CDK1.

Such a mechanism also provides an explanation for the surprising coordination of mitochondrial ROS production and CDK2 activity during the cell cycle. While we cannot exclude that mitochondrial ROS is already important for T160 phosphorylation on cyclin E-CDK2 complexes during the entry into S phase, our experiments are consistent with the idea that a limiting function of ROS for cell cycle progression lies within S phase itself (Figures 1 and 3, 4).

How does the redox state of C177 regulate KAP binding on the molecular level? The structure of KAP in association with CDK2-pT160 indicates that the interaction surfaces of both proteins are distant from C177 (Song et al. 2001), but only residues 25-198 of KAP were resolved and truncation of residues 1-34 completely abolishes its interaction with CDK2 (Yeh et al. 2003). Hence, it is conceivable that the N-terminus of KAP directly binds to CDK2 in a C177 oxidation-sensitive manner. Alternatively, and beyond the scope of our present study, we are currently investigating whether an unknown CDK2 interactor binds to C177 in its oxidized form and thereby prevents the recruitment of KAP (Figure 7K). KAP itself contains several redox-sensitive cysteine (Xiao et al. 2020) and cysteine 140 (C140), crucial for KAP activity is easily oxidized *in vitro* (Song et al. 2001). Thus, the increase of ROS during S and G2 phase might also negatively regulate KAP activity and thereby feedback on CDK2 activity. While evidence that C140 can be oxidized in living cells is still lacking our data suggest that, if existing, this alternative mechanism of CDK2 regulation is independent of CDK2 itself because mutating C177 to serine makes the CDK2-KAP interaction insensitive to the redox state of the cell (Figure 7K and S3J).

As cells become cancerous and rewire their metabolism to promote proliferation and survival, they often produce increased amounts of ROS (Reczek & Chandel 2017). Our findings provide a molecular mechanism for how a cancer cell might exploit increased ROS to drive proliferation via CDK2. They also indicate how this unique feature of CDK2 activity regulation could be exploited to develop CDK2-specific inhibitors that do not target closely related kinases, i.e. by a cysteine-reactive probe that prevents C177 oxidation but still allows KAP binding.

---

**Figure S1, related to Figure 1 and Figure 3.**
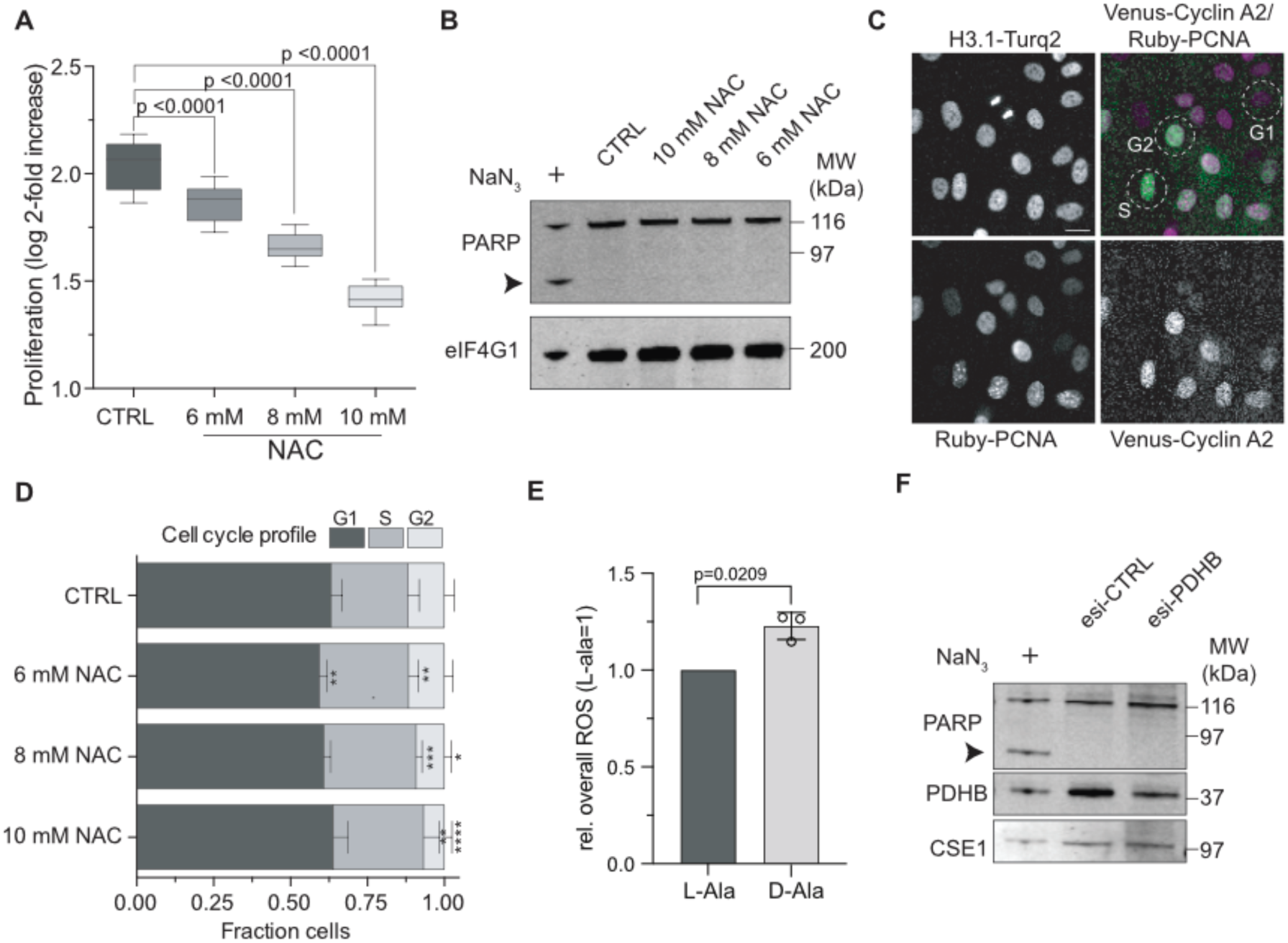
Interference with ROS slows cell proliferation and does not induce apoptosis. (A) Proliferation of RPE-1 cells in the presence of the indicated concentrations of N-acetyl-L-cysteine (NAC). Boxplots indicate the median log2-fold increase of cells during 48 hours NAC treatment. Significance according to one-way ANOVA with Dunnett’s multi comparison test (n=3, N=18). (B) Representative Western blot analysis detecting PARP cleavage in cells depleted of PDHB or treated with NAC to assess induction of apoptosis. Note PARP cleavage (arrowhead) is only visible upon treatment of cells with 50 µM sodium azide (NaN_3_), a known apoptosis inducer; detection of eIF4G1 serves as a loading control. (C) Exemplary cell cycle analyses of single cells from (A) using endogenously-tagged H3.1-Turquoise2 (H3.1-Turq2) to segment and cyclin A2-Venus and Ruby-PCNA to unambiguously classify cells. Cyclin A2-Venus negative cells are in G1 phase, cyclin A2-Venus positive cells are in S or G2 phase, and PCNA-foci identify S phase. Circles indicate examples of classified cells. Scale bar = 10 µm. (D) Stacked bars showing the mean ± cell cycle phase distribution of the cells from (A) and classified as indicated in (C). Significance according to one-way ANOVA with Holm-Sidak’s multicomparison test: **(6 mM NAC (G1) = 0.0077), **(6 mM NAC (S) = 0.0085), ***(8 mM NAC (S) = 0.0007), *(8 mM NAC (G2) = 0.0133), **(10 mM NAC (S) = 0.0046), ****(10 mM NAC (G2) = p<0.0001), (n=3, N=18). (E) Quantification of ROS detected by CellRox Deep Red in HyPer2-DAO-NES expressing RPE-1 cells as in Figure 1F in response to 0.5 mM D-alanine (D-ala) Control cells were treated either with 0.5-or 5-mM L-alanine (L-ala). Bars represent the mean ± SD. Significance according to two-tailed one-sample t-test (n=3, N=3). (F) Representative Western blot analysis assessing apoptosis by virtue of PARP cleavage in cells depleted of PDHB; detection of CSE1 serves as a loading control.

**Figure S2, related to Figure 5 and Figure 7.**
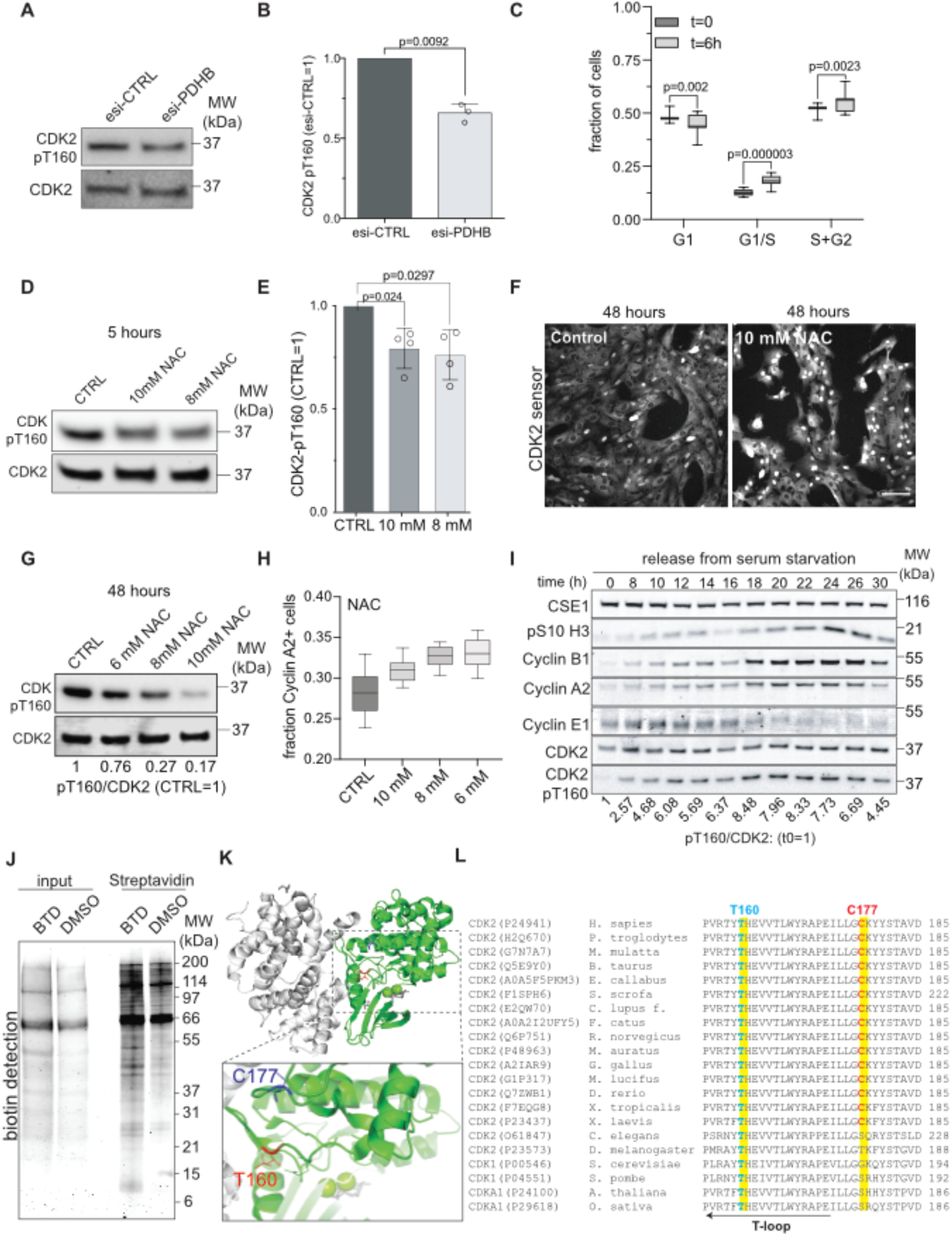
Changes of CDK2 T-loop phosphorylation during the cell cycle and in response to reducing treatments and identification of C177 oxidation. (A) Representative Western blot analysis showing T160 phosphorylation in cells depleted of PDHB for 48 hours and synchronized at the beginning of S phase with thymidine. (B) Quantification of the data shown in (A) Bars indicate the mean ± SD. Significance according to two-tailed one-sample t-test (n=3, N=3). (C) Cell cycle distribution analysis based on live imaging of Fucci hGem(1-110) and Cdt(30-120) expressing cells. G1 phase cells are Cdt1 positive but hGem negative, S phase cells are positive for both markers, G2 phase cells are positive for hGem but negative for Cdt. Boxplots indicate the median fraction of cells in the indicated cell cycle phase at the beginning and after 6 hours of glutamine starvation. Significance according to two-way ANOVA with Sidak’s multicomparison test (n=3, N=18). (D) Representative Western blot analysis showing the degree of T160 phosphorylation in S phase-synchronized cells 5 hours after treatment with the indicated concentration of NAC. (E) Quantification of the data shown in (C). Bars indicate the mean ± SD. Significance according to two-tailed one-sample t-test (n=4, N=4). (F) Representative (n=3) image showing the response of CDK2 activity sensor expressing RPE-1 cells to 48 hours of treatment with 10 mM NAC. Note, nuclear-localized sensor is indicative of CDK2 with low activity. Scale bar = 100 µM. (G) Representative Western blot analysis to (n=2) detect pT160 in RPE-1 cells treated with the indicated concentrations of NAC for 48 hours. (H) Single-cell imaging analysis of Cyclin A2-Venus expressing cells (see Figure S1C) treated as in (G). Boxplots indicate the median fraction of Cyclin A2 positive cells in response to the indicated treatments (n=3, N=18). (I) Representative Western blot analysis (n=2) showing the T-loop phosphorylation and the expression of the indicated cell cycle markers in RPE-1 cells after the release from serum starvation (24 hours). The ratio of phosphorylated T160 (pT160) to CDK2, normalized to T=0 is indicated beneath the Western blot. (J) Representative Western blot analysis of denaturing Streptavidin pull downs showing the detection of biotinylated proteins from the BTD labeling experiment presented in Figure 7B. (K) Structure of CDK2-cyclin A (PDB: 4I3Z) highlighting T160 and nearby C177. (L) Clustal Omega alignment of CDK2 kinases showing that C177 is highly conserved in vertebrates. Note, organisms in which CDK2 functions are exerted by a single CDK (i.e., in yeast or plants) do not contain a corresponding cysteine residue.

**Figure S3, related to Figure 7.**
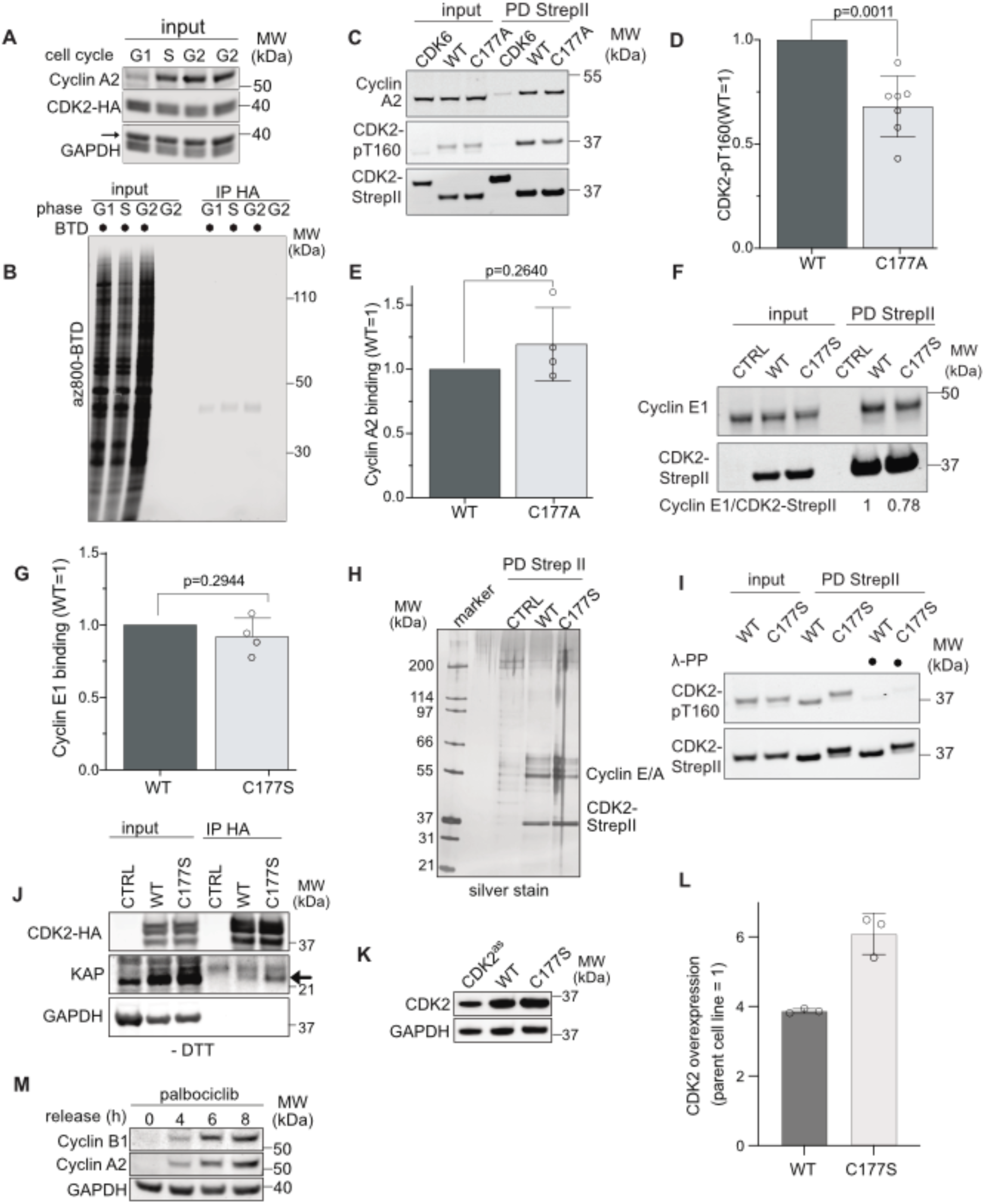
Modulating the oxidation state of cysteine 177 affects T-loop phosphorylation of CDK2 and proliferation. (A) Representative Western blot analysis of input samples of (B) and data in Figure 7D showing the differential expression of cell cycle marker cyclin A2 in G1 versus S and G2 phase samples. (B) Complete scan of the data presented in Figure 7D at a lower intensity to highlight differential BTD labeling of input samples at different stages of the cell cycle. (C) Representative Western blot analysis of StrepII pull downs using cell extracts from S-phase synchronized RPE-1 cells transiently expressing CDK2-WT-StrepII (WT), CDK2-C177A-StrepII (C177A) or CDK6-StrepII as a control. The relative degree of T160 phosphorylation (pT160) and co-precipitated cyclin A2 normalized to WT and corrected for loading is shown beneath the blot. (D-E) Quantification of pT160 (E) and cyclin A2 binding (F) from data shown as in (C). Bars represent the mean ± SD. Significance according to two-tailed one-sample t-test. (pT160: n=7, N=7; cyclin A2: n=4, N=4). (F) Representative Western blot analysis of StrepII pull downs as in (C) showing the interaction of cyclin E1 to CDK2-StrepII WT and C177S. (G) Quantification of cyclin E1 binding from data shown in (F). Bars represent the mean ± SD. Significance according to two-tailed one-sample t-test. (n=4, N=4). (H) Silver staining of Strep II pull downs of CDK2 WT and C177S from lysates of S phase synchronized cells in the absence of DTT. Note, magnetic beads-bound CDK2-StrepII from presented samples were used for the experiments shown in Figure 7H. (I) Representative Western blot analysis showing the dephosphorylation of T160 by λ-phosphatase (λ-PP) on magnetic beads-bound CDK2-StrepII WT and C177S used for CAK *in vitro* binding assays in Figure 7H. (J) Representative Western blot analysis (n=3) of CDK2-HA immuno-precipitates from asynchronous WT and C177S expressing RPE-1 cells. Note, the lysates were prepared without reducing reagents, which prevented the binding of KAP to WT but not to C177S CDK2. (K) Representative Western blot analysis of total lysates from parent CDK2^as^ cells and CDK2^as^ cells stably expressing untagged CDK2 WT or C177S. (L) Quantification of CDK2 WT and C177S expression shown in (K). Bars represent the mean and ± SD. (n=3, N=3). (M) Representative Western blot analysis (n=2) of RPE-1 from a 24-hour Palbociclib induced G1 phase arrest to establish that the 4 and 6-hour time points analyzed in Figure 7N correspond to early and mid S phase.

**Movie 1: Replication foci formation and S phase progression in esi-CTRL treated cells.** Single-cell time-lapse imaging in RPE-1 mRuby-PCNA cells showing replication foci formation every 7 minutes after at least 24 hours of esi-CTRL treatment. For better visualization of the nucleus the segmentation mask nucleus is shown. Scale bar = 20 µm.

**Movie 2: Replication foci formation and S phase progression in esi-PDHB treated cells.** Single-cell time-lapse imaging in RPE-1 mRuby-PCNA cells showing replication foci formation every 7 minutes after at least 24 hours of esi-PDHB treatment. For better visualization of the nucleus the segmentation mask nucleus is shown. Scale bar = 20 µm.

## EXPERIMENTAL PROCEDURES

### Cell culture and cell treatments

All cell lines were cultured according to standard mammalian tissue culture protocol at 37 °C in 5% CO_2_ and tested for mycoplasma contamination. For hypoxia experiments cells cultured under standard cell culture conditions were adapted to 6.3% (RPE-1) and 4% O_2_ (BJ) in Whitley H35 hypoxystation (Don Whitley Scientific) for 3-4 days before the experiment. Prior to subsequent imaging or flow cytometry analyses cells were fixed at the indicated concentrations of O_2_ in 3.7% in PFA/PBS for 15 minutes to maintain the hypoxic phenotype. hTERT RPE-1 (RRID:CVCL_4388), hTERT RPE-1 FRT/TR (RRID:CVCL_VP32), hTERT-RPE-1 CDK2^as^ (Merrick et al. 2008) and the thereof derived cell lines listed in Supplemental Table 1 were grown in DMEM/F12 (Sigma Aldrich) supplemented with 10% (v/v) FBS (Gibco), 1% (v/v) penicillin-streptomycin (Sigma Aldrich), 1% (v/v) Glutamax (Gibco), 0.5 μg/mL Amphotericin B (Sigma Aldrich) and 0.26% sodium bicarbonate (Gibco). HEK-293 (RRID:CVCL_0045) cells were grown in DMEM (Gibco) supplemented with 10% (v/v) FBS, 1% (v/v) penicillin-streptomycin, 1% (v/v) Glutamax, 0.5 μg/mL Amphotericin B. BJ fibroblasts (RRID:CVCL_3653) were grown in MEM (Sigma Aldrich) supplemented with 10% FBS, 1% (v/v) penicillin-streptomycin and 1% (v/v) Glutamax (Gibco). For BJ fibroblasts, the surface of cell culture plates was functionalized with 1% gelatin prior seeding. To synchronize RPE-1 cells at the beginning of S phase 2.5 mM or 10 mM thymidine (Sigma Aldrich) were added to the growth medium for 24 hours. To synchronize cells in G1 phase 150 nM Palbociclib (Sigma) was supplied to the medium for 24 hours. For glutamine starvation experiments cells were washed once with PBS and incubated for 6 hours in DMEM/F12 without glutamine and supplemented with 10% (v/v) dialyzed FBS (Gibco), 1% (v/v) penicillin-streptomycin, 0.5 μg/mL Amphotericin B, and 0.26% sodium bicarbonate. 500 mM stock solutions of N-acetyl-L-cysteine (NAC, Sigma Aldrich) in 7.5% sodium bicarbonate were always prepared freshly, adjusted with NaOH to pH 7.4-7.5, and added to the cell culture medium at the indicated concentrations and time. PEG-Catalase (Sigma) was dissolved in 50% glycerol/H_2_O and added to the cells at the indicated amounts. 1 M stock solutions of D- and L-alanine in H_2_O (Sigma Aldrich) were stored at -20 °C and diluted to the indicated final concentration into growth media 24 hours after cell seeding for proliferation experiments (Figure 1) or 48 hours after esi-RNA transfection (Figure 5). MitoTEMPO (Sigma) was dissolved in DMSO and added at the indicated concentrations. Endogenous CDK2 in RPE-1 CDK2^AS^ cells was inhibited by adding 10 µM 3MB-PP1 (Merck) for the duration of the experiment. To ensure normal CDK-cyclin pairing (Merrick et al. 2008) 0.5 µM 6-benzylaminopurine (6-BAP) was added during cultivation and experiments as indicated. As a positive control for apoptosis (Figure S1) 50 µM sodium azide (Sigma Aldrich) was added to RPE-1 cells for 24 hours.

### Plasmids and cell line generation

All plasmids and cell lines used in this study including vector backbones, PCR primers, templates, restriction sites, parent cell lines, inserts, methodology, and sources are listed in Supplemental Table 1 and 2. Knock in’ s of mRuby-PCNA and Histone 3.1-iRFP were created by rAAV-mediated gene targeting as described previously (Zerjatke et al. 2017; Collin et al. 2013). Stable cell lines were created by electroporation of plasmids using a Neon Transfection system (Thermo Fisher) according to manufacturer instructions. 14-21 days later single positive clones were picked after selection with 400 μg/ml neomycin (Invitrogen) or 0.5 μg/ml puromycin (Sigma Aldrich). RPE-1 cell lines ectopically expressing HyPer2-DAO-NES or HyPer2-DAO-NLS were generated by viral transduction of cells using an rAAV gene targeting system (Stratagene) and subsequent isolation of single Hyper2-fluorescent clones by cell sorting on a BD FACSAria III (BD Biosciences). All cell lines used in this study including parental cell lines, used plasmids, methods of creation, and references are listed in Supplemental Table 1. For transient transfection 8 µg of CDK2-StrepII WT or C177S plasmid were electroporated per 1 mil. cells followed by extract preparation 48 hours later (see below).

### RNA interference

esi-RNA or si-RNA oligonucleotides were delivered using RNAiMAX (Thermo Fisher Scientific) by reverse transfection according to the manufactures instructions. Briefly, transfections contained 16.5 ng esi-RNA per 96 wells, 49.5 ng esi-RNA per µ-Slide 8 Wells, or 1 µg esi-RNA per 6 well, or 50 nM si-RNAs mixed with 0.2 µl, 0.6 µl and 2 µl of RNAiMAX in OPTIMEM (Gibco), respectively. RNAi-treated cells were analyzed 48 hours after transfection, exception of long-term time-lapse imaging (Figures 3G and 4A), which was started already after 24 hours. esi-PDHB (HU-04685-1, Eupheria), esi-CTRL (5’-TGTCCCTTAAACACTCACTGGTCACGAGCGATACAATTCGCATACGGAGATAGGAGAATCGTCATACGTCGATACAGGTGCATAAAACGGCCTTCCAAGATTCGTCGATCTAATATTTTCGGGGGACGATTAATATAAATGGGTCTTCTACAAGTCTATTGATCATAGTTCTTAACGTAGGGACGTTCGTTACATGAAATAAGACTTAGTTACCACACTTCAATATTCATTTTGCCCGACCTGTCGCCAG-3’, Eupheria), si-PDHA (5’-AGGUUGUGCUAAAGGGAAA-3’, Eurofins) and si-CTRL (5’-UGGUUUACAUGUCGACUAA-3’, Dharmacon)

### Proliferation experiments and EDU detection

To determine initial and final number of cells for proliferation experiments the complete 96 well was imaged 5 hours after seeding (for esi/si-RNA experiments) or just prior to compound application and at the end of the experiment, respectively. Cell number were determined by detection and segmentation of nuclei by virtue of fluorescently tagged histone 3.1 or staining with 200 nM life cell DNA dye SIR-DNA (Spirochrome). For cell competition CDK2as cells stably expressing CDK2 WT and GFP from the same RNA (CDK2 WT_IRES2_eGFP) or CDK2 C177S and mRuby (CDK2 C177S_IRES2_mRuby) were seeded into the same 6 well in the presence of 10 µM 3MB-PP1. Cell numbers of WT and C177S cells were determined by splitting a 1000-15000 cells from the 6 well into a 96 well plate, followed by incubation with 200 nM SIR-DNA and imaging of cells 1 hour after seeding when attachment of all cells was complete. WT (GFP positive) and C177S (mRuby positive) were identified by single cell analysis nuclear based on nuclear segmentation and GFP/mRuby detection using the segmentation and filter modules of MetaXpress Custom Module Editor (Molecular Devices). The ratio of WT/C177S cells at day 0 was set to 1 and the procedure repeated at day 3 and 6. DNA replication was assessed by adding 10 µM EDU (Thermo Fisher) to cells for 45 minutes (PDHB depletion and MitoTEMPO experiments) or for the duration of the experiment (Figure 7N). Afterwards cells were fixed in 4% PFA/PBS for 15 minutes at RT, washed twice in 3% BSA/PBS and extracted for 20 minutes in 0.5% Triton X-100/PBS, followed by two 5 minutes washes in 3% BSA/PBS. A click mixture in PBS containing 4 mM CuSO_4_ (Sigma), 5 mM AF647-Picolyl-azide (Jena Biosciences) and 10 mM sodium ascorbate (Sigma) was added to cells for 1 hour at RT in the dark. After removal of the click mixture cells were washed once in 3% BSA/PBS and PBS followed by staining of DNA with 2 µM Hoechst 33342 for 30 minutes, two washes with PBS and image analysis.

### Lysate, extracts and interaction studies

For total lysates RPE-1 cells were washed once in PBS followed by the addition of 1x NuPAGE LDS Sample buffer (Novex) including 100 mM DTT were briefly sonicated and boiled at 95°C for 5 minutes. For CDK2 pT160 detection RPE-1 cells were trypsinized, washed in PBS and resuspended in extraction buffer P (30 mM Tris pH 7.5, 0.25% NP-40, 2.5 mM MgCl_2_, 175 mM NaCl, 10% glycerol, 1 mM DTT, cOmplete protease inhibitors (Merck), phosSTOP phosphatase inhibitors (Merck)), incubated for 20 minutes on ice, and centrifuged at 13,000g for 15 minutes at 4°C. Cleared extracts were mixed 3:1 (v/v) with 3x NuPAGE LDS/DTT Sample buffer (Sigma Aldrich) and boiled at 95°C for 5 minutes. For StrepII pull downs RPE-1 cells were lysed with extraction buffer S (50 mM Tris pH 8, 150 mM NaCl, 2.5 mM MgCl_2_, 5% glycerol, 1% Triton X-100, cOmplete protease inhibitors (Merck) and incubated for 20 minutes on ice. Extracts were cleared (13,000g, 15 minutes, 4 °C) and added to extraction buffer-equilibrated MagStrep “type3” XT beads (IBT), and incubated for 30 minutes on a rotating wheel at 4 °C. To assess binding of endogenous KAP to CDK2-StrepII-WT, living cells were incubated with 5 mM DTT at 37 °C for 10 minutes prior to extraction in buffer S (-/+ 20 mM DTT). For analyses of CDK2 T160 phosphorylation, cyclin A2 binding, cyclin E1 binding, and endogenous KAP binding beads were washed 3x in extraction buffer S and precipitates were eluted in NuPAGE LDS/DTT sample buffer at 95 °C for 5 minutes.

For binding assays with recombinant KAP (Sigma Aldrich, Cat. No. SRP5175) or CAK (Thermo Fisher, Cat. No. PV3868) CDK2-WT-StrepII and CDK2-C177S-StrepII immobilized on MagStrep “type3” XT beads were washed 1x in extraction buffer S -/+ 10 mM DTT. For CAK binding assays CDK2 was dephosphorylated by washing beads once in 1x λ-phosphatase buffer (NEB) supplied with 1 mM MnCl_2_ (NEB) and incubated with 400 units of λ phosphatase (NEB, Cat. No. PO753L) for 30 minutes at 4 °C. Subsequently, CDK2 WT or C177S beads were incubated with or without 10 mM DTT in extraction buffer S including 0.1 µg/µl bovine serum albumin (BSA) for 30 minutes at 4 °C. KAP binding assays were performed for 30 minutes on a rotating wheel at 4 °C with 60 ng KAP supplied to binding buffer K (50 mM Tris pH 6.8, 150 mM NaCl, 0.1 µg/µl BSA) -/+ 1 mM DTT. CAK binding assays were performed for 1 hour on a rotating wheel at 4 °C with 125 ng of CAK diluted in binding buffer C (8 mM MOPS/NaOH pH 7, 0.2 mM EDTA, 150 mM NaCl) -/+ 1 mM DTT. After binding, beads were washed twice with extraction buffer S and eluted with in NuPAGE LDS/DTT sample buffer for 5 min at 95 °C. For binding of endogenous KAP to CDK2-WT-StrepII, living cells were incubated with 5 mM DTT at 37 °C for 10 minutes before extract preparation in buffer S -/+ 20 mM DTT. Staining of CDK2-StrepII pull downs was performed using the ProteoSilver stain kit (Sigma) according to the manufactures instructions.

### BTD labeling

RPE-1 cells were grown to 80-90% confluency in 15 cm dishes (Greiner Bio-One), washed once in PBS, and labeled for sulfenic acids with 1 mM of the sulfenic acid-reactive probe BTD (Gupta et al. 2017) dissolved in DMSO and added to the growth medium for 30 minutes at 37 °C (0,5% final concentration DMSO). Subsequently, cells were washed twice with PBS and fixed in PBS/1% formaldehyde (FA) supplemented with 10 mM iodoacetamide (IAA) for 15 minutes. After further alkylation in PBS/10 mM IAA for 15 minutes, cells were reduced with 40 mM DTT/PBS for 30 minutes, washed once with PBS, permeabilized with 90% -20 °C-cold methanol for 15 minutes at -20 °C, washed once with PBS, and treated with 15 µg/mL benzonase per 15 cm dish for 15 minutes to digest DNA and RNA. After a further wash in PBS, cells were blocked with PBS/5% BSA for 1 hour, washed once in PBS, and incubated with 0.5 nmol picolyl-azide PEG4-biotin in 7.5 ml PBS (Jena Bioscience) per dish. Clicking of Biotin-azide to BTD-alkene was catalyzed by a 2x concentrated click mix to reach a final concentration of 1 mM CuSO_4_, 0.1 mM Tris-[(1-benzyl-1*H*-1,2,3-triazol-4-yl)-methyl]-amin (Sigma Aldrich), and 1 mM sodium ascorbate. After clicking for 1 hour at RT, cells were washed once with PBS, thrice for 10 minutes in PBS/0.1% Tween 20/10 mM EDTA, once with PBS, and scraped into 1 ml PBS containing 2% SDS/5 mM DTT. Then, samples were boiled at 95 °C for 45 minutes to reverse the FA crosslink, cleared for 15 minutes at 13,000g, supernatants were diluted 6 times in PBS, added to PBS-equilibrated magnetic streptavidin beads (Pierce), and incubated on a wheel over night at 4 °C. Subsequently, unbound material was removed, beads were washed 4x in 4 M urea/0.5% SDS/25 mM HEPES, 4x in PBS/0.5% SDS and transferred to a clean microfuge tube. After two further PBS washes bound proteins were eluted by NuPAGE LDS/DTT sample buffer supplemented with 2.5 mM biotin/100 mM DTT at 95 °C for 15 minutes. To investigate the cell cycle stage dependent oxidation of CDK2, RPE-1 cells expressing CDK2-HA WT were grown to 50% confluency and serum starved for 24 hours. According to Figure S2I cells were labelled at 8, 18 and 22 hours representing G1, S and G2 phases for 30minutes at 37 °C in media containing 1% FBS to minimize BTD probe scavenging by FBS. Subsequently, cells were washed twice with PBS, lysed in 50mM Tris pH 8, 400mM NaCl, 5% Glycerol, 1% SDS, supplemented with fresh f.c. 10mM TCEP and protease inhibitors (Roche) for 15 minutes at RT followed by alkylation (f.c. 40mM IAA) for 15 minutes at RT in the dark, stored at -80°C until all time points were collected, sonicated to shear DNA and cleared at 13.000g. In every replicate 1 – 2 mg BTD-labeled lysate was clicked to Az800 (AzDye 800 – Click Chemistry tools) as described above. Afterwards, proteins by Chloroform/Methanol precipitation to remove click reagents and protein pellets were resuspended in 20mM Tris pH 8, 0.5M Urea, 0.5% SDS. Lysates were sonicated and diluted 1:7 with buffer A containing 20mM Tris pH 8, 120mMlNaCl, 1% Triton and incubated protein G Dynabeads (Thermo Fisher) coupled to HA antibodies for 1 hour at 4°C. Then, beads were washed once with buffer A supplied with 120 mM NaCl and thrice with buffer A with 400mM NaCl. CDK2-HA was eluted by boiling beads for 10 minutes at 65°C in NuPage LDS sample buffer, followed by transfer to a new tube and addition of 100 mM DTT and analysis by SDS-PAGE.

### SDS-PAGE and Western blot analyses

Proteins were separated by SDS-PAGE using Bis-Tris 4–12% gradient gels in MES or MOPS buffers (Lifetech) in a Mini Blot Tank (Thermo Fisher). Western blot analyses were performed using a wet transfer Criterion Blotter (BioRad) in MOPS/20% ethanol transfer buffer using Immobilon-FL PVDF membranes (Millipore). Membranes were blocked for 1 hour at RT in 5 % dry milk (Roth) or 5 % soy protein isolate (Vitasyg, for CDK2-pT160 detection) prepared in PBS/0.2% Tween 20 (Life scientific). Primary antibodies were added overnight at 4 °C, followed by 3x washing in PBS/0.2% Tween 20 and incubation with secondary antibodies for 1 hour at RT. For quantitative detection, fluorescently-labelled secondary antibodies and a near-infrared scanning system (Odyssey, LICOR) were used. Alternatively, detection was performed with horseradish peroxidase (HRP)-conjugated antibodies and Luminata Forte Western HRP Substrate (Millipore) or Super Signal West Femto Maximum Sensitivity Substrate (Thermo Fisher) on an ImageQuant LAS4000 system (Amersham Biosciences). All primary and secondary antibodies including dilutions, product numbers and manufactures are listed in Supplemental Table 3.

### ROS detection

For flow cytometry analyses on an LSR Fortessa FACS (BD Bioscience) analyzer 1×10^6^ asynchronously growing cells were labeled for 30 minutes with 5 μg/ml Hoechst 33342 (Sigma), trypsinized, washed in PBS, and transferred to CO_2_-independent L15 media (Life Technologies) supplemented with 10% FBS and 1% (v/v) penicillin-streptomycin (Sigma Aldrich), 1% (v/v) Glutamax (Gibco), 0.5 μg/mL Amphotericin B (Sigma Aldrich) and 0.26% sodium bicarbonate (Gibco). Then 5 μg/ml Hoechst 33342 and 5 µM CellRox Deep Red (Invitrogen) and/or MitoSox Red (Invitrogen) were added for 30 minutes or 100 nM MitoTracker Green (Invitrogen) for 10 minutes. ROS labelling of adherent cells was performed in living RPE-1 cells expressing endogenously-tagged histone 3.1-Turquoise2, Ruby-PCNA, cyclinA2-Venus using the reagents, concentrations and labelling times indicated above, followed by a brief wash in PBS to remove the excess of free dye and incubating cells in imaging DMEM (see below).

### ADP/ATP ratio

The ADP/ATP ratio was determined using an ADP/ATP ratio assay kit (Sigma Aldrich) 48 hours after PDHB depletion in a 96 well plate format. Measurements were performed according manufacturer instructions on a Glomax Luminometer (Promega).

### Microscopy

Automated microscopy was performed on an ImageXpress Micro XLS wide-field screening microscope (Molecular Devices) equipped with 10x, 0.5 NA and 20x, 0.7 NA, Plan Apo air objectives (Nikon) and laser-based autofocus. Excitation and detection were done by a Spectra X light engine (Lumencor) and sCMOS (Andor) camera with the indicated filters, respectively: DAPI (Ex: 377/50; Dic: 344-404 / 415-570; Em:447/60); CFP (Ex: 438/24; Dic: 426-450/467-600; Em: 483/32); GFP (Ex: 472/30; Dic: 442-488 / 502-730; Em: 520/35), YFP (Ex: 513/17; Dic: 488-512 / 528-625; Em: 542/27), TexasRed (Ex: 575/25 Dic: 530-585 / 601-800; Em: 624/40) and Cy5 (Ex: 628/40; Dic: 594-651 / 669-726; Em: 692/40). During experiments cells were maintained in a stage incubator at 37 °C in a humidified atmosphere of 5% CO_2_. All cells were grown in 96 well plastic bottom plates (μclear, Greiner Bio-One). For long-term time-lapse microscopy and ROS imaging the growth media was exchanged to an auto-fluorescence-reduced imaging DMEM(Schmitz & Gerlich 2009) supplemented with 10% (v/v) FBS, 1% (v/v) penicillin-streptomycin, 1% Glutamax, 0.5 µg/mL Amphotericin B and 0.26% sodium bicarbonate. Long-term time-lapse microscopy was performed by taking images using a 10x objective in 7 minutes intervals for 48 hours. To determine the initial and final number of cells for proliferation experiments the complete 96 well was imaged 5 hours after seeding (for esi/si-RNA experiments) or just prior to compound application and at the end of the experiment, respectively. Microscopy of cells stained with ROS-sensitive dyes was performed by acquiring maximum image projection using a 20x objective. For ratio imaging of Hyper2-DAO-NLS and Hyper2-DAO-NES cells on the ImageXpress Micro XLS the following filter sets were used: Ex: (438/24) with Em: (542/27) and Ex: (513/17) with Em: (542/27). For CDK2 sensor imaging in cells grown under hypoxia experiments cell were fixed in 3.7% PFA/PBS prior exposure to atmospheric O_2_ during imaging

For ratio imaging to assess differences in ROS content between esi-CTRL and esi-PDHB treatments in Hyper 7 expressing cells a Delta Vision Core wide-field deconvolution fluorescence microscope equipped with a CoolSNAP HQ2/HQ2-ICX285 camera (Applied Precision Inc.) and a UPlanSApo 40X/NA 0.95 lens (Olympus) was used. Cells were imaged using the following filters: Ex: (381-401) with Em: (500-523) and Ex: (464-492) with Em: (500-523).

### Image and data analysis

Image analyses of single time points were performed in MetaXpress 6 (Molecular Devices) using customized image analysis pipelines. Briefly, images where flat-field and background corrected using a tophat filter, nuclei were segmented based on DNA or Ruby-PCNA labeling to create masks for fluorescence extraction. To determine the initial and final number of cells in proliferation experiments the complete 96 well was imaged 5 hours after seeding (for esi/si-RNA experiments) or prior to compound application, respectively, and again at the end of the experiment. Detection of PCNA replication foci was performed using the nuclear speckle plugin of the MetaXpress Custom Module Editor (Molecular Devices). Fucci analysis was performed by segmenting all nuclei based on SIR-DNA staining of nuclei and by merging the nuclear intensities of hGem(1-110) and Cdt(30-120) to a common segmentation mask. hGem(1-110) and Cdt(30-120) positive cells were assigned by minimum threshold filtering and the cell cycle distribution was indicated as a fraction of the sum of hGem(1-110) or Cdt(30-120) positive cells (Figure S2C). Time-lapse analysis of single cells was performed as previously described in detail (Zerjatke et al. 2017). Briefly, all images were background corrected using flat-field correction, nuclei were segmented based on histone 3.1 labelling using intensity thresholding and subsequent Watershed filtering. Time-lapse single-cell tracking was performed with a nearest-neighbor approach and subsequent manual track curation using TraCurate (Wagner et al. 2021). Cell cycle phases were classified according to PCNA mean intensity and distribution as described in (Zerjatke et al. 2017). Nuclear CDK2 levels were quantified as mean intensities based on the histone segmentation masks. Cytoplasmic CDK2 levels were quantified as the mean intensities in two cap regions adjacent to the poles of each cell nuclei. Image analysis and quantification were performed with Mathematica 12.1 (Wolfram Research Inc.). Thresholds for CDK2^LOW^/CDK2^High^ cells were determined by histogram analyses of ratios of nuclear and cytoplasmic CDK2 to identify the CDK2^Low^ peak (Spencer et al. 2013). Ratio image analyses were performed in Fiji (Schindelin et al. 2012) using the Ratio Plus plugin as described in (Kardash et al. 2011). Ratio image analyses of HyPer7 images were performed using semi-automated Fiji macro. Briefly, the images were background corrected (rolling ball = 50), smoothened and a mask was defined based on the HyPer7 Ex: (464-492)/Em: (500-523) image and binarized. Then, images from both excitations were multiplied with the mask to remove background, converted to 32 bit, and the ratio image was calculated by dividing Hyper 7 signal from (Ex: 464-492)/(Em: 500-523) by (Ex: 381-401)/Em: (500-523). Flow cytometry analyses were performed in FlowJo (Becton, Dickinson and Company). Forward scatter values were used as a relative proxy for cell size.

### Statistical methods

Data normalization was performed in Microsoft Excel (Microsoft), statistical analysis and graph presentation in Prism 6-9 (GraphPad) using the statistical tests indicated in the legends of each figure. All data are representative of at least three independent repeats if not otherwise stated. The notation n refers to the number of independently performed experiments, the notation N to the number of data points used for statistical analyses and data presentation. A p-value lower than 0.05 was considered statistically significant and individual p values are indicated within each figure or figure legend. Bar charts indicate the mean ± SD and show all data points for N<10. Box plots show the median and 25th-75^th^ percentiles, whiskers the 5th-95^th^ percentiles. For quantitative Western blot analyses by near-infrared fluorescence detection (Li-Cor) band intensities were divided by the intensity of the loading control or by the intensity of total CDK2 levels (pT160 blotting) to corrected for unequal sample loading. Time-lapse data of single-cell ratio imaging were smoothing with the 4 neighbors on each size and a 6th order polynomial using Prism 6 (Graph Pad). For flow cytometry analyses the intensities of individual cell cycle phases were normalized by the sum of intensities of all cell cycle phases to correct for different overall staining intensities in between experiments. No randomization or blinding was performed in this study.

### Data availability

Generated plasmids, cell lines and scripts used for data analyses are available from the corresponding author upon request.

## Acknowledgments

We thank André Nadler for the synthesis of BTD, Vsevolod V. Belousov for sharing Hyper7 before publication, Sabrina Spencer for sharing CDK2 activity sensor plasmids, Robert Fisher for sharing RPE-1 CDK2^as^ cells and experimental suggestions, Kate Carroll and André Nadler for sharing BTD, Alex Radzisheuskaya for help with cell competition assays, Jon Pines for critical reading of the manuscript, Thuy Dung Vu and Marina Cuenca for help with single-cell tracking and ratio analyses, respectively, and members of the Mansfeld laboratory for their technical help and discussions. J.M. receives funding from the European Research Council (ERC) under the European Union’s Horizon 2020 research and innovation program (grant agreement no. 680042). D.K., K.J. and J.V. are members of the Biomedicine and Bioengineering (DIGS-BB) Ph.D. program. We acknowledge support by the Light Microscopy and flow cytometry facilities of the Center for Molecular and Cellular Bioengineering (CMCB) of the TU Dresden and the Institute of Cancer Research UK (ICR).

## Author contributions

Conceptualization, J.M, D.K.; Methodology, D.K., K.J., J.V., T.Z., I.G. and J.M.; Investigation, D.K., K.J., J.V., J.K.L., and J.M.; Writing—Original draft, D.K., K.J., and J.M.; Funding acquisition, I.G. and J.M.; Supervision, I.G. and J.M.

